# Organotypic tissue architecture is a requisite for predictive outcomes of perfluorocarbon exposure in airway models

**DOI:** 10.1101/2024.10.29.620983

**Authors:** Shawn W H Liu, Filbert O Christone, Ryan D Quarrington, Eugene Roscioli

## Abstract

**Aims:** Perfluorocarbons (PFCs) are inert, oxygen-rich fluids with applications in neonatal and adult liquid ventilation and potential for innovations related to hypoxic environments, such as under water and for space. Despite promising clinical applications, a laboratory model to comprehensively test PFC implications and predict human airway responses is lacking. We hypothesise that an organotypic airway epithelial cell (AEC) model is needed assess PFC impacts and provide predictive outcomes to inform the *in vivo* scenario that conventional submerged cultures cannot replicate. This study evaluated PFC exposure in two *in vitro* systems: a traditional submerged model using 16HBE14o- cells and an air-liquid interface (ALI) model using hSABCi and primary nasal AECs, cultured on transwells to mimic the human airway epithelium.

**Methods:** PFC exposures were conducted at 2, 8, and 24 hours (submerged model) and extended to 72 hours (ALI model). We tracked gross morphological changes via microscopy, quantified apoptosis and autophagy markers through protein biochemistry, and assessed epithelial permeability using trans-epithelial electrical resistance (TEER) and tight junction protein abundance. Statistical analyses included at least three biological replicates per condition.

**Results:** In the submerged 16HBE14o- model, PFC exposure led to significant apoptotic changes by 24 hours, with marked autophagic disruption. Nutrient deprivation, confirmed by starvation experiments, was a key driver of cytotoxicity due to media/PFC phase separation. Conversely, hSABCi cells in the ALI model remained viable, with no apoptosis or autophagic disruption over the exposure periods (P > 0.05 vs. control). Similarly, primary nasal AECs showed consistent viability and stability in autophagic and apoptotic markers, indicating a more accurate representation of in vivo conditions. TEER measurements and tight junction protein levels in the ALI model suggested PFC did not compromise epithelial integrity (P > 0.05 vs. control). PBS exposure, included as a liquid control, underscored baseline sensitivity in the primary nAEC model, evidenced by SQSTM1 upregulation and pronounced barrier dysfunction over 24-72 hours (P < 0.05–0.001). These findings underscore the unique biocompatibility of PFC in maintaining cellular integrity, in contrast to the disruptive effects observed with PBS.

**Conclusion:** This is the first study to describe PFC exposure in an organotypic airway model. Results indicate that the ALI model more accurately preserves airway epithelial integrity during PFC exposure than submerged models, which are limited by nutrient depletion effects. The findings support the use of ALI cultures to replicate the human airway architecture for evaluating PFC’s biological effects and offer a platform for preclinical applications in respiratory medicine. This organotypic approach may inform future therapeutic and hypoxia-related interventions and contributes significantly to the field by providing a viable model for understanding PFC interactions with airway epithelial cells. We are now trialling PFC emulsions that also contain azithromycin and steroid therapies for extended therapeutic applications.

**Infographic:** 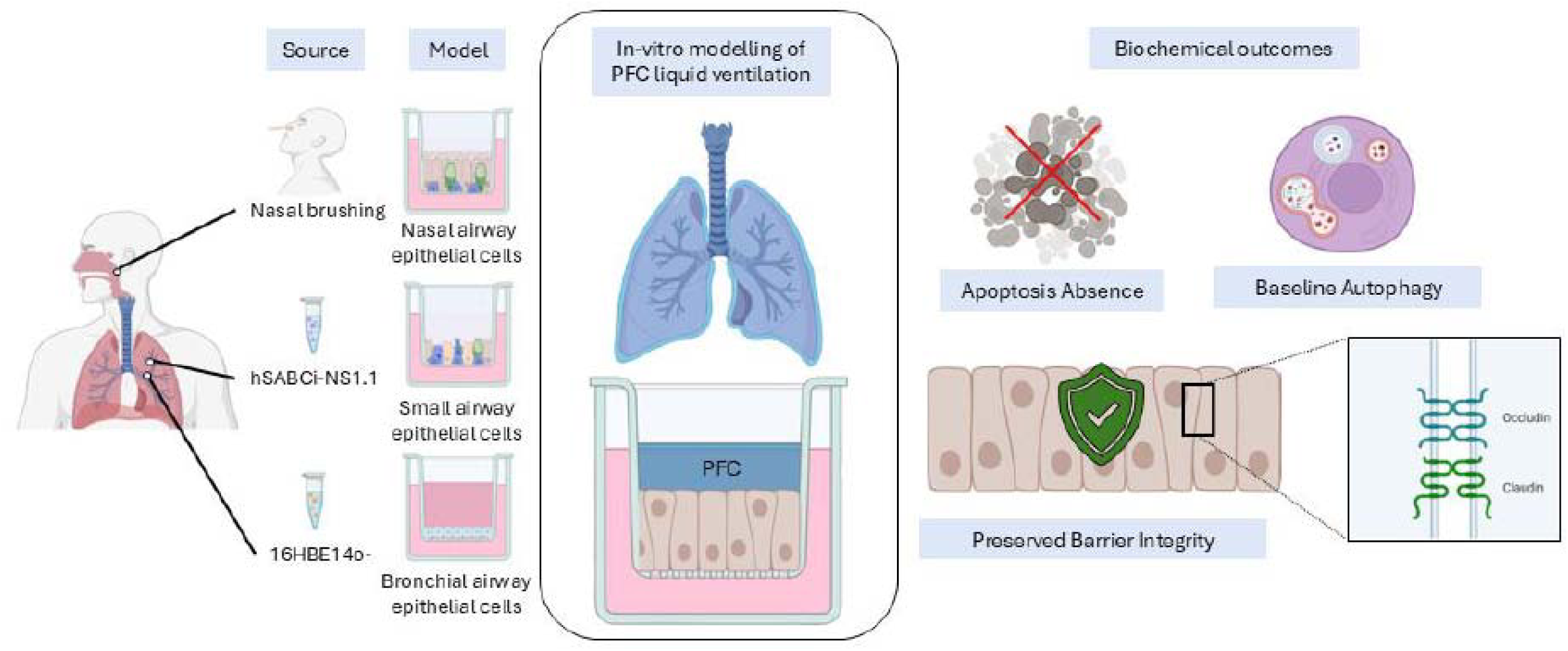

## Introduction

While terrestrial animals exchange oxygen and carbon dioxide from the air, it is also possible for the lungs to exchange respiratory gases from a select family of biologically inert liquid fluids – perfluorocarbons (PFC). PFC liquid breathing is used in respiratory care and is largely unknown to the public and most respiratory (or otherwise) scientists. This leaves ample innovation space, with few models that can pursue new applications. In current medicine, liquid breathing with PFC liquid ventilation has been researched in the past decades as an alternative to conventional mechanical ventilation (CMV) to provide respiratory assistance to patients with severe lung conditions, particularly premature infants with underdeveloped lungs. However, many conceivable innovative approaches for PFC and liquid breathing are yet to be investigated, such as human endeavours in low-oxygen environments, operating in pressures that cause decompression sickness underwater or enhancing human ability to withstand G-force in space.

The research performed herein sets the platform for a needed and essential method to assess PCFs in the controlled environment of the laboratory by applying state-of-the-art airway models to this exciting technological approach for enhanced pulmonary function. It is hypothesised that PFC can be examined in highly representative models of the human airways to identify the toxicity, viability, and function of the airway epithelium in the context of liquid breathing.

It is foreseeable that the work to develop this model will provide a much needed incentive for the broader scientific community to bridge the knowledge gap that currently limits medical pulmonology and future directions that extend the endeavours of human exploration in otherwise inhospitable environments..

### Human airway epithelium

The human airway epithelium is a crucial component of the respiratory system, functioning as a physical barrier to the external air environment and an integral part of the innate immune system. The airway epithelium lines the respiratory tract, extending from the protective epithelium of the nasal cavity down to thin alveolar walls [1], where gas is transmitted to the primary internal respiratory tissues at the alveoli (respectively)[2]. Furthermore, the airway epithelium possesses multifaceted functions depending on the region (upper or lower airway). It comprises various specialised cell types, including ciliated (enable coordinated particle clearance), goblet (produce a highly complex and immunologically essential mucus lining), and basal cells (progenitors of all airway cells). Ciliated and goblet cells are responsible for mucociliary functions, where goblet cells secrete mucus airway debris and unwanted pollutants, and ciliated cells beat in coordinated movement to remove the mucus [2]. Together, they enable mucociliary function, which is needed for minute-by-minute viability of the whole organism, as evidenced by the devastating consequences of cystic fibrosis [3]. Basal cells are essential cell types with differentiation capabilities to maintain and repair the epithelium following injury or cellular turnover [4]. Furthermore, the epithelium also forms tight junctions between cells and is usually apically (air facing) located in polarised airway epithelial cells (AEC); the tight junction is part of the epithelial barrier function and is among the first molecular gatekeepers that govern paracellular permeability [5].

Therefore, maintaining airway epithelium function is essential for respiratory function. Respiratory distress syndrome (RDS) is an umbrella term for a lung condition that causes low blood oxygen; it is a common pathology in dysfunctional airway epithelium, such as in patients with lung injury or premature infants with insufficient lung development [6]. Liquid ventilation has emerged as an innovative solution in the past few decades to facilitate oxygenation and recovery in RDS conditions [7] and understanding how the PFC liquid interacts with airway epithelium from a cellular and molecular level is crucial in optimising and advancing its medical applications. As the airway epithelium is the immediate interface between the environment and the underlying pulmonary tissues it protects, a model that assesses this extensive immuno-respiratory barrier is the most pertinent way to predict outcomes for PFC exposure in humans.

### Perfluorocarbon

Perfluorocarbon (PFC) is an organic compound consisting of fluorocarbon chains where all hydrogens have been replaced by fluorine [8]. Fluorine is the most electronegative (reactive) element, forming a strong polar covalent bond with carbon. This, combined with the molecular symmetry of PFC, the electron cloud from fluorine atoms, shields the carbon backbone and protects the fluorocarbon bonds from external influences [8], resulting in a highly inert and stable PFC. It is also important to note that PFCs used in medical applications are generally considered safe and not to be confused with other fluorinated chemical of per- and polyfluoroalkyl compounds (PFAS), which are associated with health and environmental concerns [9].

Perfluorodecalin (PFD, CCCFCC) and perfluorooctyl bromide (PFOB, CCFCCBr) are the PFC compounds currently used in liquid ventilation [10]. PFOB is more commonly used in liquid ventilation and is also utilised as a contrasting agent during MRI and ultrasound imaging [11]. Both PFD and PFOB possess exceptional oxygen-carrying capacity, approximately 20 times that of saline [8, 10], and their low surface tension allows efficient coverage across the lung surfaces to facilitate oxygen delivery [8, 12]. To date, there have been no other chemicals that can serve this function, making PFCs a highly valuable means to explore liquid ventilation.

### History and development of liquid ventilation

The first documented instance of perfusing mammalian lungs with liquid for medical purposes occurred after the First World War. Researchers discovered that canine lungs could tolerate saline lavage procedures without adverse effects [13]. However, the most impactful liquid breathing studies were performed in the 1960s [14, 15]. Dr Klystra demonstrated that mammalian lungs (mice) could survive for hours submerged under a pressurised and oxygenised saline liquid [14]. Meanwhile, Leland Clark and Frank Gollan demonstrated PFC as a superior liquid breathing medium compared to silicon oil, which exhibited extreme toxicity [15]. All subjects died shortly after weaning off silicon oil respiration, whereas subjects that respired PFC survived for weeks.

Moreover, further liquid breathing studies involved the first human liquid ventilation study of oxygenated saline conducted in American Naval Research [16]. The study demonstrated the feasibility of human liquid ventilation with effective gas exchange, a short recovery period and minimal discomfort for anesthetised subjects during the procedure. However, saline solution was determined to be an inadequate breathing medium as subjects experienced respiratory acidosis due to ineffective CO_2_ removal during procedures [14, 16]. Thus, it was apparent that liquid ventilation using biocompatible PFC that can facilitate both O_2_ and CO_2_ exchange is the best-suited way to move forward with liquid breathing research.

### Current applications of liquid ventilation

Dr Thomas Shaffer and his group invested most of their resources into liquid ventilation for neonatal care by utilising premature lamb models to improve liquid ventilation methodologies for neonatal application [17–19]. This leads to the first FDA-approved liquid ventilation study in three near-death neonates [20] with severe respiratory distress syndrome (RDS). Despite improved oxygenations and improvement in lung distensibility, all infants succumbed after 19 hours of liquid ventilation, and deaths were determined to be due to disease severity before ventilation. Nonetheless, this demonstrated the potential of liquid ventilation to transition premature infants to extrauterine life. Subsequent studies further explored different ventilation techniques, notably a study in 1996 that performed partial liquid ventilation (PLV) with PFOB in 13 premature infants with RDS [21]. The study demonstrated marked improvements in lung function and oxygenation that led to the survival of 8 infants to a corrected gestational age of 36 weeks. It is noteworthy that some of the surviving infants are still alive today. These outcomes results solidified the foundation for the current use of liquid ventilation in neonates. However, PFC technology is not widely used medically for reasons that likely concern the need for more supporting research in this area.

Consequently, despite the successes in animal studies and neonatal applications, clinical trials of PLC in adult RDS patients have yielded mixed results. A pivotal study in 2006 compared gas ventilation (CMV) and liquid ventilation (PLV) in approximately 300 patients with acute RDS [22]. The study found that CMV patients had more ventilator-free days compared to the PLV group and that the PLC group also experienced higher frequencies of hypoxic and hypotensive episodes, as well as pneumothoraxes. This 2006 study highlighted the limitations of PLV in adults, and adult clinical trials have since slowed down. The current focus of PFC liquid ventilation studies, which largely involve animal subjects, is on improving and refining methodologies that can be translated to clinical use. These include using PFC thermal conductivity to induce hypothermia and provide cardio- or neuroprotection after cardiac/respiratory failure [23–25] or to increase safety and cardiopulmonary stability during liquid ventilation in premature animal models [26, 27].

### *In vitro* studies of PFC liquid ventilation and gaps in understanding

Despite in-vivo animal studies and human trials over the years providing valuable insight into systemic effects and administration methods of liquid ventilation, there needs to be more in-vitro studies that investigate intricate interactions between PFC and the respiratory epithelium at a cellular and molecular level. Indeed, this is likely due to the technical expertise needed to model the human airways in an organotypic manner (e.g. [28]), coupled with relaxed ethical standards prior to the Declaration of Helsinki (1964) and the need to operate within the guidelines of subsequent institutional research review boards (late 20th century). Understanding the effects of PFC exposure on airway epithelial cells (AECs) is crucial to applying sensitive technology around liquid ventilation experiments that produce objective outcome measures predictive of the situation in vivo. This approach facilitates the identification of effective treatment regimens, particularly those not amenable to traditional clinical research, and informs further investigations in animal models [29].

Current available in vitro studies mainly investigate the anti-inflammatory effects of PFC observed during clinical trials. For example, studies explored whether PFOB creates a physical barrier that dampens inflammatory response during liquid ventilation. A study found that PFOB prevents immune cell adhesion once it diffuses through alveolar space [30] and inhibits direct cellular interactions with neutrophils. Another study found that PFOB decreased cytokine secretion in TNF-α stimulated A549 alveolar cells [31]. Furthermore, PFC also affects AEC barrier function, as observed in [32], where an increase in transepithelial electrical resistance (TEER) may have protected Calu-3 cells from hyperoxia-related injury. In contrast, PFOB causes actin cytoskeleton remodelling in A549 cells and weakens the cells’ adhesion ability [33].

Most PFC in vitro studies utilise traditional in vitro models that lack the human airway’s complex architecture and cellular characteristics and often rely on cancer-derived cell lines, like A549 or Calu-3, to assess PFC-epithelial interactions. Such models cannot inform the normal airway epithelium and are frequently used as they are easy to apply, widely available and require entry skill level. Further, these models typically lack diverse cell types that express airway surface liquid that is characteristic of their epithelial origin (see, for example [34, 35],). Moreover, mucociliary function, including mucus production and coordinated cilia beating, are vital components of the airway epithelium that have yet to be researched. Outcomes from those mentioned above in vitro studies using commercially available cancer-based models exhibit altered cellular processes and responses. In contrast, primary human cells in the ALI offer a bone fide epithelial phenotype and lend to personalised outcomes for distinct people and diseases (e.g. cystic fibrosis and asthma).

There is an apparent disconnect between previous models that examine PFC exposures and the need for outcomes that can provide information that translates to real-world applications for health and innovation. This is more so for findings to inform the clinic and pathology for liquid ventilation applied to patients with various lung conditions. Therefore, this thesis develops an organotypic model of human airway epithelium using a recently developed small airway cell line (hSABCi- NS1.1) and primary nasal AECs. The foreground outcomes of airway viability and function after in vitro PFC exposure are investigated. The core aim was to provide objective outcomes to inform this aforementioned research gap, provide a physiologically relevant investigation of PFC-induced responses in the human airway, and set a foundation for innovative investigations to identify extended applications for this technology.

## Methods

### Cell Line and General Tissue Culture Method

#### Human Bronchial Epithelial Cell Line (16HBE14o-)

16HBE14o- cell line was bought from Sigma-Aldrich and kept in liquid nitrogen until use. The cell line originated from a 1-year-old male heart-lung patient and immortalised with origin-defective SV40 plasmid [36]. 16HHBE is widely used for in-vitro studies particularly cystic fibrosis and ion transport studies due to upregulation of cystic fibrosis transmembrane conductance regulator (CFTR) .

#### 16HBE14o- Culture Method

The 16HBE cells were thawed from liquid nitrogen and propagated in tissue culture flask, and maintained with weekly passaging and media change every 2 days. The culture medium used was Dulbecco’s Modified Eagle Medium (DMEM; Sigma Aldrich) supplied with 10% foetal calf serum (FCS; Scientifix), 1% L-glutamine and 1% penicillin-streptomycin (pen/strep; both ThermoFisher Scientific). Incubator is set at 37 °C with 5% CO_2_ atmosphere.

Prior to experiments in submerged system, 16HBE cells were trypsinised (Lonza), resuspended in fresh DMEM, and 5 x 10^5^ cells per well were seeded onto a 12-well plate. For experiment in transwell model, 1x10^5^ cells were seeded onto collagen (STEMCELL Technologies) coated transwell (0.4 µm pores, 6.5 mm diameter; Sigma-Aldrich), with both compartments filled with media. Air-liquid interface (ALI) was introduced (air-lifted) on day 7, and exposure experiment was conducted 3 days post airlift.

#### Human Small Airway Basal Cells immortalized-Nonsmoker 1.1 (hSABCi- NS1.1)

The hSABCi cell line was originally generated in Prof. Crystal’s laboratory at Cornell University in New York [37] and was transferred to our laboratory with collaboration with Matthew Gartner from University of Melbourne at passage 52 and kept in liquid nitrogen until use. This is a recently developed cell line derived from a healthy 50-year-old African American male and immortalised through retrovirus transduction of telomerase reverse transcriptase gene [37]. Additionally, hSABCi is a progenitor cell line that retains differentiation capabilities that are region-specific characteristic of the small airway epithelium.

#### hSABCi Culture Method

The hSABCi cells were thawed and suspended in Ex-plus media and cell count was performed to confirm the cell viability before propagating the cells on a collagen (STEMCELL) coated flask.

Culture medium was PneumaCult Ex-plus Medium (STEMCELL), 1 x 10^5^ cells were seeded onto 6.5 mm Transwell inserts (Sigma-Aldrich) pre-coated with collagen. Following cell expansion, the cultures were air-lifted, and the media was replaced with complete PneumaCult ALI Basal Medium (STEMCELL) supplied with 1% pen/strep to promote differentiation at an air-liquid interface (ALI). Experiments were conducted on hSABCi that were airlifted for at least 25 days.

#### Primary Human Nasal Airway Epithelial Cell Line (nAEC)

The primary human nasal airway epithelial cell air–liquid interface culture model was generated as previously described [1]. Nasal samples were obtained from one adult volunteer (45-years-old) in compliance with the Declaration of Helsinki and written consent of the volunteer. Ethics approval was granted by the Central Adelaide Local Health Network Human Research Ethics Committee. The use of one primary nasal sample number was sufficient to ensure the hSABCi model was applicable to the experiments described here.

Briefly, volunteer underwent nasal brushing, and brushes were rinsed in complete Ex-plus media to obtain basal AEC. Rinsed sample was centrifuged for 300g for 5 mins at 10 °C, and pellet was resuspended in Ex-plus media before propagated onto a collagen coated tissue culture flask. Media change was performed daily until nAEC cell reached confluent. Subsequently, cells were trypsinised, neutralised (Lonza) and seeded at 1x10^5^ cells in collagen coated 6.5mm transwell insert. Media was replaced every day until day 5 where nAEC cells were airlifted and complete ALI Basal media was used in the basal compartment to signal differentiation. Media was changed every second to third day and mucus clearance was performed every 4-5 days with 5 minutes apical PBS (ThermoFisher) incubation, followed by secondary PBS wash for 10 seconds.

### Perfluorocarbons and Exposure Method

#### Perfluorocarbons (PFC)

The PFCs used in this thesis are perfluorodecalin (PFD; Sigma-Aldrich) and perflubron/ perfluorooctyl bromide (PFOB; ThermoFisher). These are the PFC primarily used in liquid ventilation.

#### PFCs Exposure in Transwell

150 µl of PFC was applied to the apical compartment at the start of exposure. PFC were left in the apical compartment and untouched except for TEER modules where the PFC will be replaced with HEPES for the period of TEER measurement and refilled back. Empty wells around the transwell are filled with PBS to increase humidity and avoid PFC evaporation during exposure period. For exposure period longer than 24 hours, apical PFC level was checked every day and refilled if evaporation was observed.

#### Microscopy

Routine light microscopy was performed to assessed cell morphology during expansion and differentiation as well as throughout various experimental modules. Observations were conducted using a Nikon Eclipse Ts2 inverted microscope. Images were captured using TrueChrome 4k Pro camera, operated using the Mosaic 2.4.1 software (Tuscen). Camera calibration was performed at each objective lens magnification (4x, 10x, 20x, 40x) using the etching of Neubauer improved haemocytometer (Parul Marienfield). Image scales are generated based on these calibrations.

#### Transepithelial Electrical Resistance (TEER) Measurement

Evaluation of airway epithelial barrier function was performed by measuring transepithelial electrical resistance (TEER). Cell culture plates were maintained at 37°C in the biosafety cabinet using a heating platform (Leica Microsystems). Prior to measurements, treatments/exposures were aspirated from the apical compartment and filled with 150 µl of HEPES buffered solution (Lonza).

STX4 EVOM electrodes connected to an EVOM 2 electrical impedance reader (both World Precision Instruments) were used to measure electrical resistance in ohms (Ω). Electrodes were sterilised in 70% ethanol and rinsed twice with HEPES solution to avoid contamination between wells. The shorter electrode was positioned in the apical compartment, while the longer electrode was positioned in the basal compartment. TEER readings were recorded once the number stabilised. Raw resistance values are presented as Ω/cm^2^ in this thesis.

During TEER measurement of **16HBE14o-** cells. An inconsistency in the apical fluid used for different sample groups was introduced due to unfamiliarity with the technique: For the untreated (NT) samples, DMEM media was used during TEER measurement at the time points of 2 and 6 hours, as opposed to PBS that was used for all other exposure groups. This inconsistency was identified and rectified during the 24-hour endpoint, where all groups used PBS as apical fluid during TEER measurement. The potential impact of this technical inconsistency will be discussed in chapter 3 and 4.

### Sample Extractions

#### Basal Media and Apical Washes

Basal media remained unchanged during PFC exposure period and conditioned media are extracted and transferred into Eppendorf tube at the end of experiment. For experiment modules in the transwell system, HEPES solution are applied apically and allowed to incubate for at least 5 minutes. To avoid signal dilution, the HEPES washes did not exceed 300 µl. Following media extraction and apical wash, both samples were stored at -80°C until further analysis.

#### Cell lysis and protein extraction

Cells were lysed using 65 µl of M-PER Mammalian Protein Extraction Reagent (MPER) supplemented with Halt protease inhibitors (both ThermoFisher) for submerged cultures, and 45 µl for transwell cultures. Lysis was performed on ice by directly applying the M-PER/Halt solution to the cultures, followed by immediate scraping of the cells from the well-plates and transwells. Samples were frozen in -20 °C freezer before thawing and spun at 13500g for 10 minutes to remove cellular debris. The supernatant was transferred to new Eppendorf tube is stored for later used.

#### Western Blot

Protein concentrations were determined using the Pierce BCA Protein Assay Kit (Thermo Fisher), with minor modifications to the manufacturer’s protocol. Briefly, a bovine serum albumin (BSA) standard curve was generated using a 1:2 dilution series (provided with the kit). Absorbance value of protein samples suspended in M-PER was measured at 562 nm, and protein concentrations were calculated based on the standard curve.

For gel electrophoresis, 10 µg of each protein sample was prepared with 3.75 µl of 4x LDS Loading Buffer, 1.5 µl of 10x Sample Reducing Agent (both Thermo Fisher), and deionised water to a final volume of 15 µl. The samples were boiled for at least 10 minutes at 72°C and loaded onto a 1 mm thick NuPAGE 4–12% Bis-Tris Mini Protein Gels (Thermo Fisher). Electrophoresis was performed for 2 hours at 100V in Thermo Fisher Mini Gel Tank or until the dye front migrated to the bottom of the gel. Proteins were transferred to a 0.2 µm nitrocellulose membrane using a Trans-Blot Turbo Transfer System (both BioRad)

Before antibody probing (Table 1 shows primary antibodies), membrane was cut in accordance with molecular weight of the protein being probed. Blocking buffer was either 5% skim milk or 5% BSA in Tris-buffered saline (ThermoFisher) with 0.1% Tween-20 (Sigma-Aldrich; TBS-T) and membrane was blocked for 1 hour at room temperature. Primary antibody incubation in the same blocking buffer was performed overnight at 4°C on an orbital shaker. Following primary antibody incubation, membrane was washed 4 times with TBS-T for 1 hour with 15 minutes interval each wash. Secondary antibodies (Table 2) were diluted in only 5% skim milk and incubation was performed for 1 hour at room temperature. The same washing procedure with TBS-T was performed for 1 hour. Chemiluminescent imaging was performed on LAS-3000 Luminescent Image Analyzer (Fujifilm) and protein densitometry was performed using Multi Gauge software (V3.1; Fujifilm).

**Table 1:** Western Blot Primary Antibodies.

| Processes / Pathways | Antibody | Catalogue number & Company |
| --- | --- | --- |
| Cytoskeletal | Mouse anti- β-Actin | #A1978 Sigma-Aldrich Co. |
| Autophagy-related | Rabbit anti Sequestosome 1 | #5114 Cell Signalling Technology |
|  | Rabbit anti LC3 I/II | #4108 Cell Signalling Technology |
| Apoptosis-related | Rabbit anti PARP | #9542 Cell Signalling Technology |
| Tight Junctions | Mouse anti Claudin-1 | #37-4900 Thermo Fisher Scientific |
|  | Rabbit anti Occludin-1 | #71-1500 Thermo Fisher Scientific |
|  | Mouse anti ZO-1 | #33-9100 Thermo Fisher Scientific |

**Table 2:** Western Blot Secondary Antibodies.

| Antibody | Catalogue number & Company |
| --- | --- |
| Anti Mouse IgG Horseradish Peroxidase-conjugated Antibody | #HAF007 R&D Systems |
| Anti Rabbit IgG Horseradish Peroxidase-conjugated Antibody | #HAF008 R&D Systems |

#### Enzyme-linked immunosorbent assay (ELISA)

Apical washes and media in basal compartment of transwell culture were collected and used to assess IL-6 and IL-8 secretion using uncoated ELISA kits (Thermo Fisher) at endpoint of PFCs exposure for hSABCi and nAEC.

#### Statistical Analysis

Statistical analysis was performed using GraphPad Prism version 10.1.0. Student’s t-test was applied to compare two groups of data; ANOVA were applied to discover significance for more than two data sets, and multiple comparisons test comparing experimental group to untreated group were also performed. Significant outcomes were assigned to an alpha value below 0.05.

## Results

### Conventional submerged airway epithelial cell models exposed to PFC exhibit protracted cytotoxic effect

#### Microscopic observations of PFC-exposed 16HBE14o- exhibit changes consistent with apoptosis

There were no observable morphological distinctions observed in the 16HBE14o- (hereafter referred to as 16HBE) model for the PFD or PFOB 2-hour treatment interval, compared to the untreated reference (media-only exposure; Figure 1A). 16HBE cells in all groups displayed homogenous cobblestone morphology consistent with terminally differentiated AECs expressing tight connections between cells typical of a normal monolayer. After 5 hours, the PFC-exposed groups remained healthy looking, although some signs of blebbing were appearing (Figure 1B), which may indicate early signs of cellular stress.

**Figure 1.**
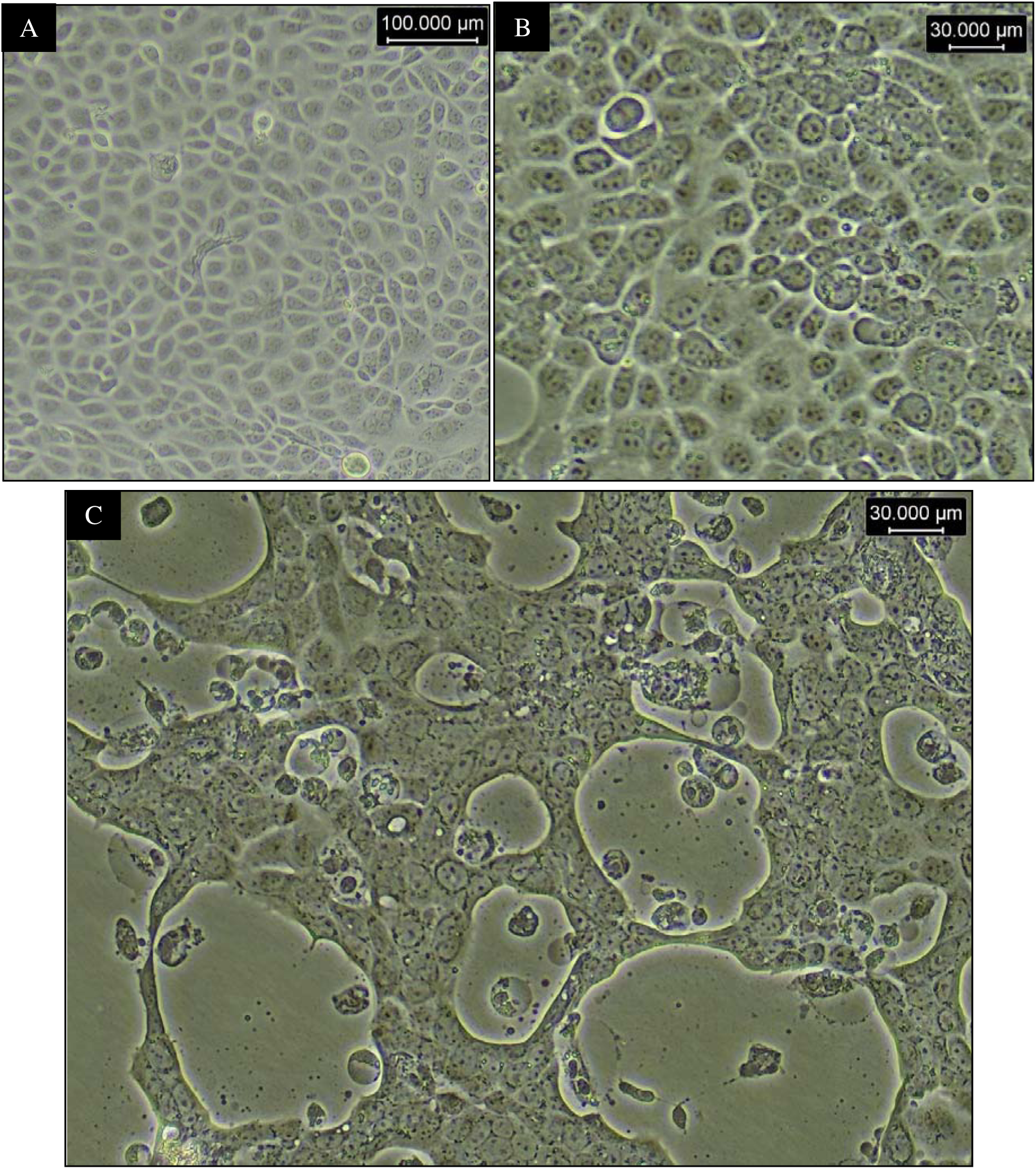
16HBE14o- under brightfield microscope during 24-hour PFC exposure. 16HBE14o- cells were exposed to perfluorodecalin (PFD) and perfluorooctyl bromide (PFOB) for 24 hours. (A) Untreated 16HBE 2 hours into experiment (10x objective lens) (B) PFOB group 5 hours into experiment (20x objective lens) (C) PFOB group 24 hours into experiment (20x objective lens)

There were clearly more discernible morphological changes between the PFC-treated and untreated control at the 24-hour interval that indicate cellular stress. The 16HBE monolayer was characterised by widespread disruptions (Figure 1C) where including zones of no cell growth “holes” observed within the monolater and cellular rounding (vs polygonal topologic morphology) combined with blebbing morphology, indicate apoptosis occurrence via cell detachment. The cause of this phenotype change was likely due to nutrient deprivation as a results of phase separation between the media and PFC due difference in specific gravity (PFC specific gravity is approximately 1.8 – 2.0 vs water). This was examined in terms of starvation-related apoptosis

#### Time-dependent cleavage of PARP (a signature of apoptosis) was observed in PFC-exposed 16HBE cells

Western blot (Figure 2A) to detect PARP protein revealed an apparent increase of cleaved PARP protein over time, with the highest blot intensity observed in the exposed group at 24 hours. Quantification of cleaved PARP blot intensities (normalised to the β-actin signal; Figure 2B) showed a similar pattern, and two-way ANOVA analysis revealed PFC exposures (P =0.0146) and exposure time (P = 0.0166), both were significant for the increase of cleaved PARP. Post hoc tests (Dunnett’s test) did not reveal any statistically significant pairwise differences due to limited sample size (n = 3 per group). However, the visual differences in cleaved PARP blot signals between untreated control and PFC groups provide compelling evidence of apoptosis occurrence.

**Figure 2.**
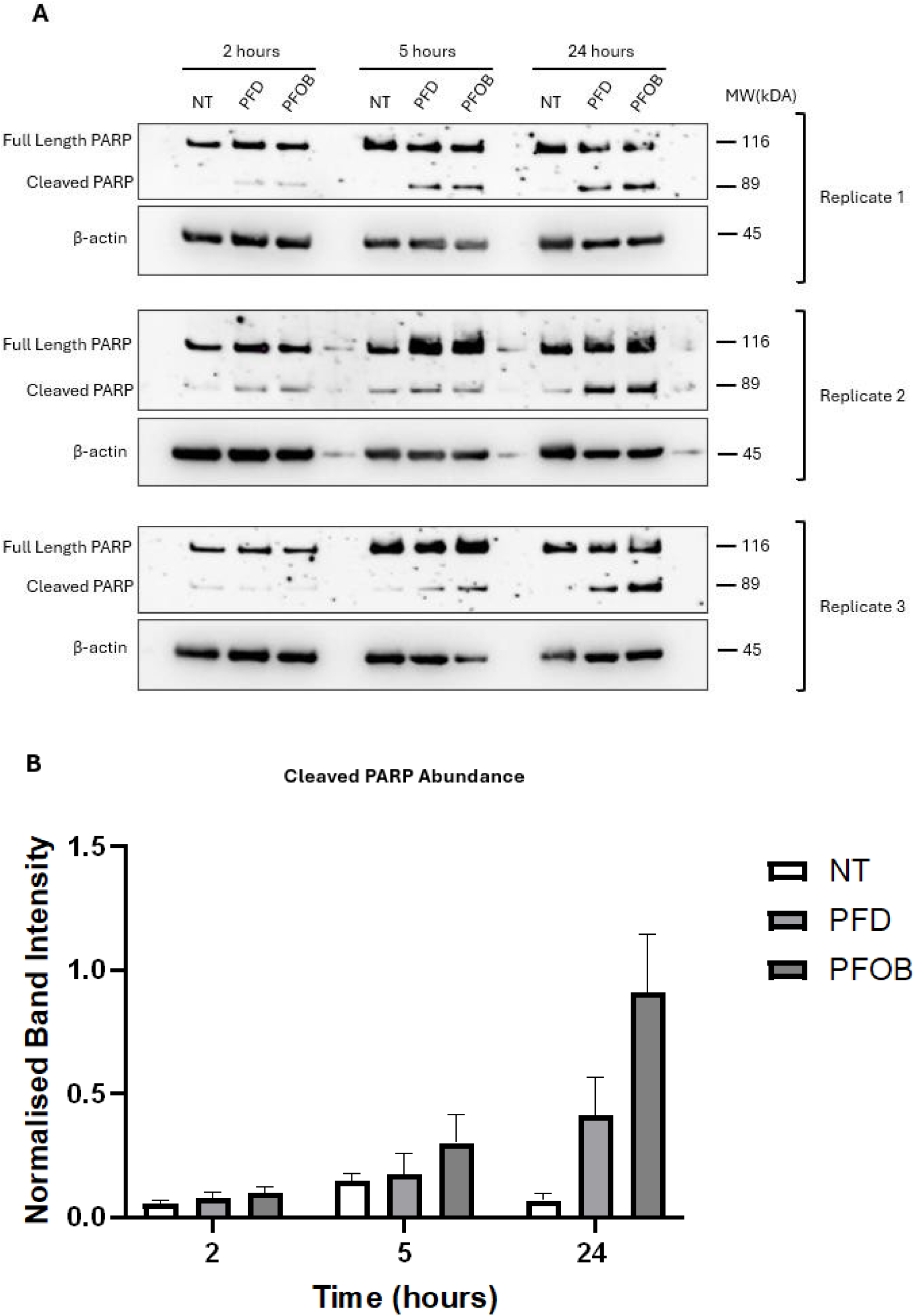
A submerged model of PFC-exposed 16HBE14o- cell demonstrated cleavage of the essential DNA repair enzyme PARP: a diagnostic molecular sign for apoptosis induction. (A) 16HBE14o- cells were exposed to perfluorodecalin (PFD) and perfluorooctyl bromide (PFOB) for 2, 5 and 24 hours, and PARP protein quantified via western blot. Membranes were probed for full length and cleaved PARP with β-actin as the loading control. Elevation in cleaved PARP was found to be positively associated with PFC exposure and therefore increased apoptotic activity. Three independent experimental replicates are presented. (B) Cleaved PARP blots were quantified and normalised to the β-actin loading control. Data are presented as bar graphs representing mean ± SEM (n = 3). Two-way ANOVA analysis revealed a significant effect of exposure time (P = 0.0166) and PFC exposure (0.0146) on PARP cleavage. These factors were also significant (0.0282), indicating the effect of exposure on PARP cleavage varied with exposure duration. However, post hoc test revealed no significant differences between PFC-exposed group and untreated group at all experiment timepoints (P>0.05 for all comparisons), which could be due to insufficient sample size.

Hence, herein valuable information was identified showing a submerged model is not possible for PFC due to immiscible media and PFC. The likely caused a starvation response (which is later assessed). To overcome this modelling limitation, subsequent cultures applied the air-liquid interface architecture which by design, are more applicable to AEC with air apically and media in the basal aspect of the cell.

#### Outcomes for PFC exposure in a submerged AEC model closely resemble nutrient deprivation in 16HBE cells

We induced a starvation response in 16HBE by decreasing (5%) or removing (0%) FCS concentration from DMEM and assessed apoptosis outcomes after 24 hours. The highest PARP cleavage signal was observed during the 24-hour time point from the last module. Therefore, the same time point was used in this module to capture the culmination of the apoptotic signal. An addition of apoptosis-positive control, doxorubicin (1 µM; known to produce a classical caspase-cascade leading to PARP cleavage [38, 39]), was used as a positive reference PARP cleavage. Figure 3A shows that complete depletion of FCS from 16HBE model induced PAPR cleavage comparable to doxorubicin control from western blot. Quantitative analysis revealed no significant differences between the starvation groups (5% FCS: P = 0.6248; 0% FCS: P = 0.3446) and the untreated group, despite an apparent increase in PAPR cleavage. This observation is similar to **3.1.2** where apoptosis occurred, but the cleaved PARP protein differences were nonsignificant between PFC groups and untreated references (could be due to low sample sizes). These similar outcomes suggest the cells exposed to PFC were indeed deprived of essential biosynthetic intermediates and nutrients, rather a direct consequence of the PFC(s). This is in line with neonatal children being exposed to PFC, and not suffering a mortal outcome due to protracted airway epithelial loss.

**Figure 3.**
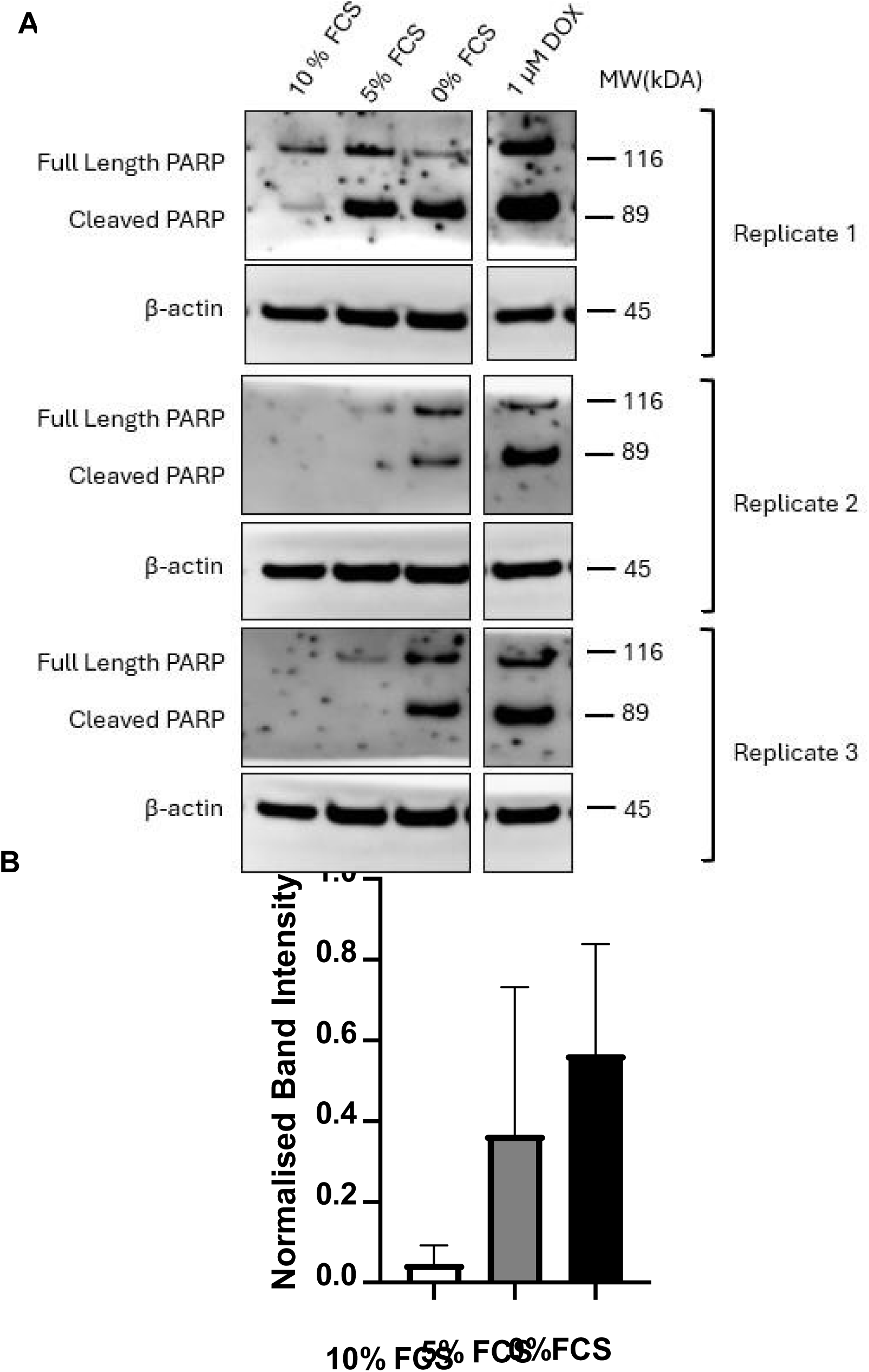
Starved 16HBE cells exhibit similar PARP cleavage pattern as PFC-exposed 16HBE in submerged model. (A) Foetal calf serum (FCS) concentration was reduced (5%) or removed (0%) in DMEM to induce a starvation response in 16HBE1. 1 µM doxorubicin (DOX) served as positive control and β-actin as loading control. 16HBE cells in all conditions were incubated for 24 hours, and proteins were extracted at the endpoint. Blots were probed against PARP protein and β-actin as loading control. Blots were edited with DOX adjacent to starvation groups for PARP cleavage comparison (B) Quantification of 16HBE cleaved PARP protein from different DMEM FCS concentrations. Data are presented as bar graphs representing mean ± SEM (n = 3 independent experiments). The data show a concentration-dependent starvation response as indicated by cleaved PARP protein abundance.

Taken together, the outcomes here support the requirement of an in vitro system that applies an air-liquid tissue engineering approach. Hence, the modelling system was revised by considering PFC’s inherent physiochemical properties (high density and hydrophobicity) and implementing a transwell-based system in subsequent experiments. Transwell technology provides an organotypic system that replicates the situation in the mammalian airways, and is predicted to solve the issues of a submerged model of the airway epithelium.

### A transwell model replicates the airway epithelial architecture and supports the production of outcome measures predictive of the scenario *in vivo*

#### 16HBEs retain epithelial integrity during PFC exposure with the transwell system

The transwells model consists of two compartments, 1. A basal reservoir for media that perfuses through 0.4 um pores of the transwell membrane that the cells are cultured on, and 2. An apical reservoir that allows the cells to be exposed to the air. This allows for the separation of PFC exposure (apically applied) and media delivery. This architecture overcomes the limitation of submerged model, and instead provides an air-liquid system. Hence where the PFC cannot interfere with the media (they are separated by the cells on the transwell. Hence, both media and PFC incident the cells basally and apically, respectively, allowing simultaneous feeding and exposure of the16HBE cells. Note: the Untreated group in scenario is 16HBE exposed to air apically, vs PFC.

Compared to submerged cultures, visualising 16HBE cells in the transwell using light microscopy is more challenging (Figure 4). The cells appeared translucent, making it difficult to observe aspects of the cytoplasmic content. However, the cobblestone-like structure and characteristic of the epithelial monolayer where tight connections join adjacent cells is still apparent. After 24-hours PFC exposure, minimal cell detachment and disruption of epithelial barrier/integrity was observed and 16HBE appeared similar in morphology to PFC-exposed vs the untreated group (Figure 4).

**Figure 4.**
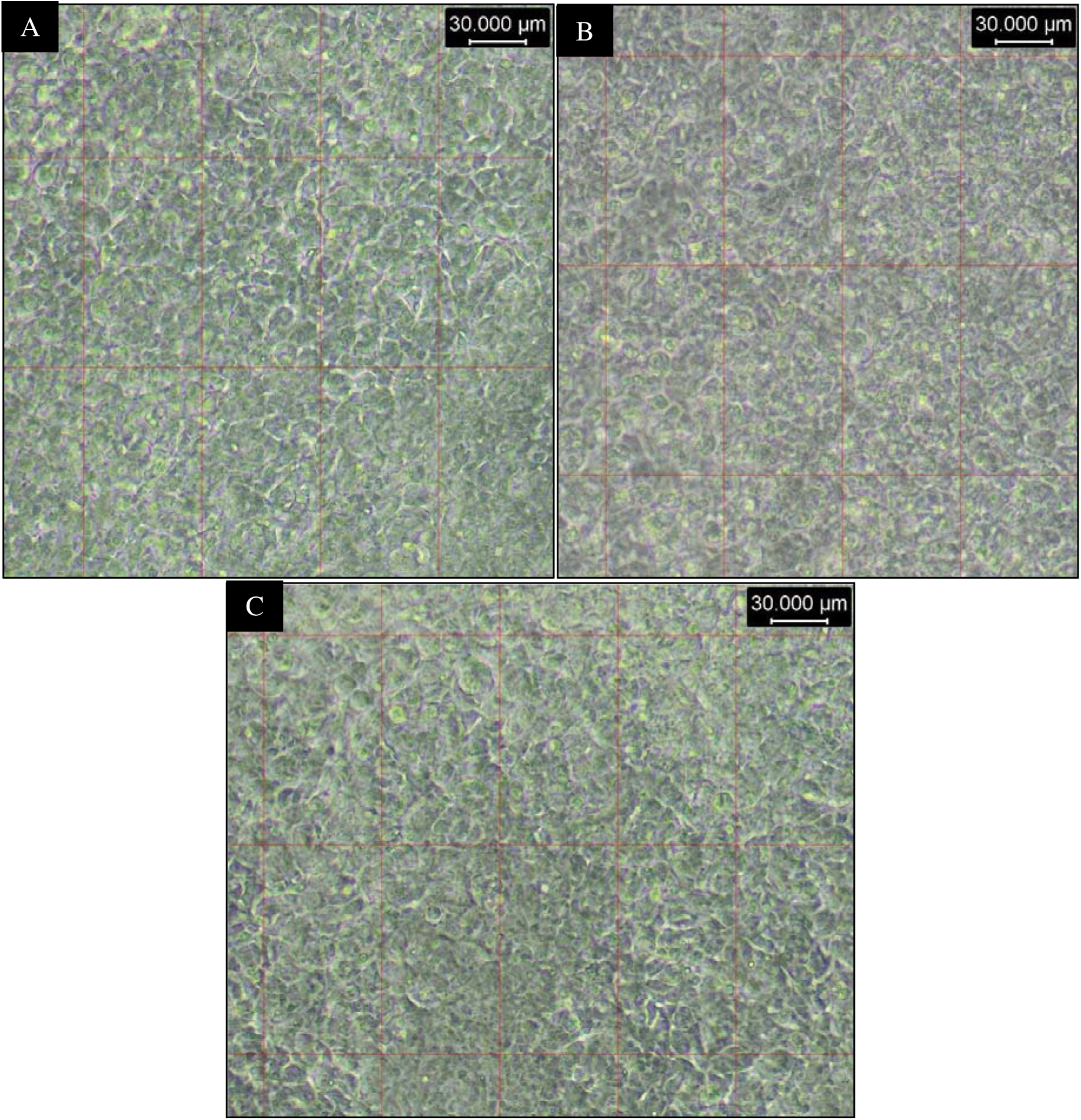
The morphology of 16HBE14o- grown on transwell and exposed to perfluorocarbons. (A) Untread/media exposed 16HBE (B) Perfluorooctyl bromide exposed 16HBE (C) Perfluorodecalin exposed 16HBE All pictures were taken at endpoint of experiment (after 24 hours), and at 20x objective lens.

#### Western blotting PARP and autophagy markers of 16HBE in transwell

To assess 16HBE cellular integrity/viability after PFC exposure, western blot analysis of PARP cleavage and autophagy markers was performed; this is an indicator of two central outcomes unambiguously related to, cellular stress viability. Apart from the PFOB group in replicate 1, all PFC-exposed 16HBE cells exhibited higher amount of cleaved PARP compared to the untreated group in Figure 5A. However, quantitative analysis revealed the differences in PARP cleavage between the PFC-exposed and untreated groups were not statistically significant (P>0.05) and were less pronounced than those observed in Figure 5A. Similarly, analysis of autophagy markers (SEQ & LC3BI/II) confirmed comparable protein abundance between untreated and PFC-exposed groups (Figure 5B), with a consistent pattern observed between replicates. Quantification of the blot outcomes (Figure 5B) confirmed these observations, but the differences were not statistically significant (P>0.05). Nevertheless, the increased level of autophagy markers indicates a mild degree of autophagy upregulation, suggesting that PFC exposure may induce cellular stress despite its inert nature, or the lack of air (vs the control) has a yet to be described effect that modulates autophagy.

**Figure 5.**
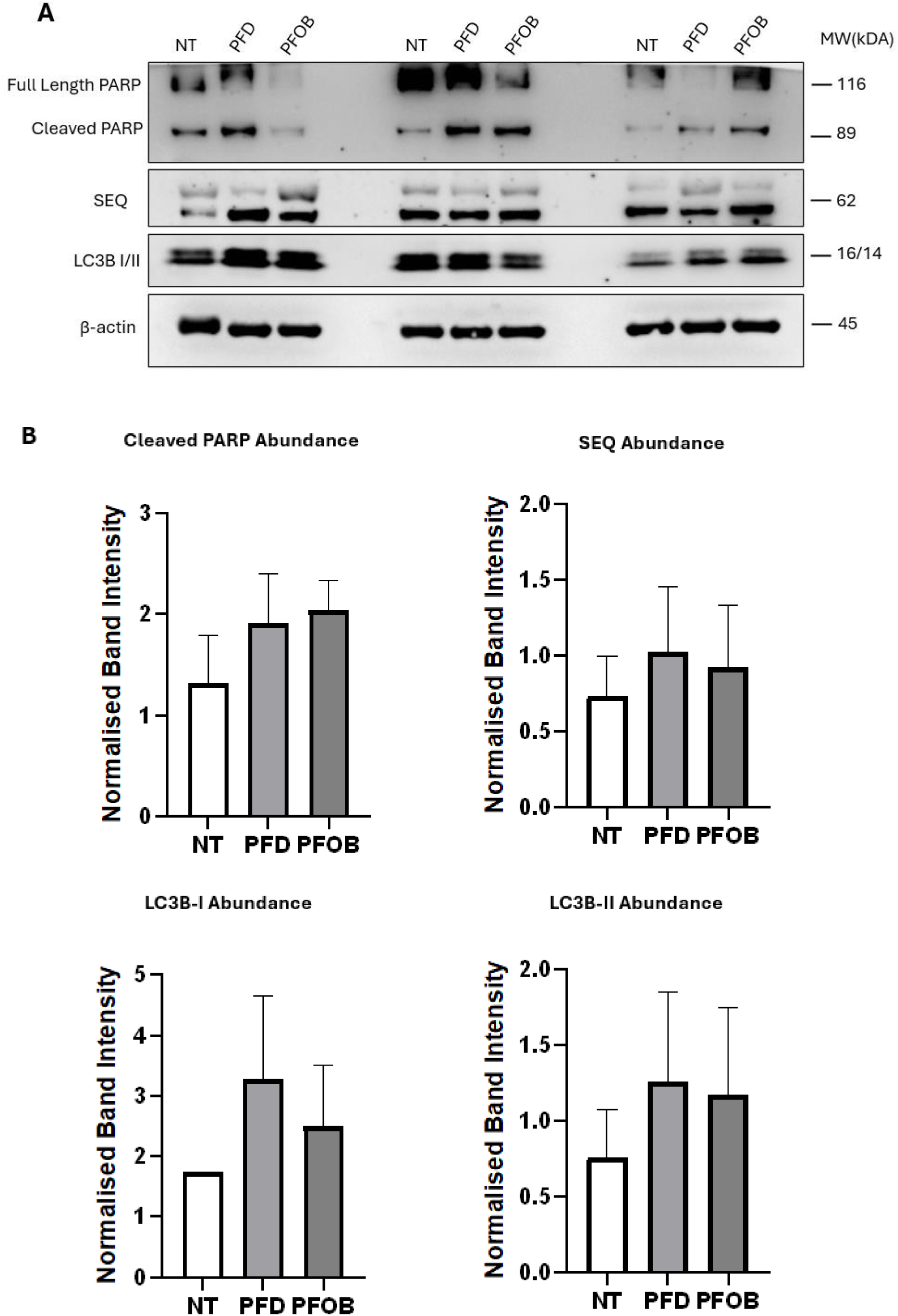
Perfluorocarbons have limited apoptotic and autophagy inducing effect in 16HBE14o- cells grown in transwell. (A) Western blot analysis of 16HBE14o- cells after 24 hours exposure of PFD (perfluorodecalin), and PFOB (Perfluorooctyl bromide) in transwell model. Sample groups from three replicates were arranged with untreated (NT), PFD, and PFOB conditions grouped adjacently for direct comparison. Blots were probed for full length and cleaved PARP (apoptosis marker), SQSTM1 (SEQ), LC3B-I/II (both autophagy markers), with β-actin as the loading control. (B) Quantification of blot intensities normalised to β-actin. Data are presented as bar graphs, with each representing mean ± SEM (n = 3). No significant differences (P>0.05) were observed between the no-exposure group and the perfluorocarbons groups

Given that 16HBE cells do not express cilia and lack a defined mucosa layer even when cultured at an ALI, further experiments were conducted using a recently developed small airway epithelial cell line, designated hSABCi-NS1.1. hSABCi are progenitor cells that can differentiate and grow to a pseudostratified epithelium that closely resembles the human small airway. The cell line’s airway surface liquid production allows for the assessment of exposure (here PFC), more predictive of the situations encountered by mammalian bronchial epithelia.

#### PFC exposure elicited an increase in the ionic conductance of the paracellular pathway, exhibited by an increase in TEER values

The transwell model allowed for the assessment of 16HBE cell barrier integrity via TEER measurement. Due to initial variations in the apical fluid used during TEER measurements (DMEM for the untreated group and PBS for all other groups), comparisons were restricted to the 24-hour time point, where all measurements were enabled by the application of PBS during the assessment interval. A control group exposed to PBS apically (i.e. for the exposure – a non-PFC liquid control) was also included in this module to assess the effect of apical liquid on 16HBE cells. TEER data for other time points are included in the appendix.

At 24 hours (Figure 6), untreated 16HBE exhibited the highest TEER value of 2551 Ω/cm^2^, the PBS group displayed a lower TEER value of 1800 Ω/cm^2^, while both PFD and PFOB produced values below 1000 Ω/cm2, One-way ANOVA analysis revealed a highly significant effect of exposure condition TEER readings (P<0.0001). Post hoc analysis demonstrated highly significant differences between the PFC-exposed groups and the untreated group (P<0.0001), and a less pronounced significant difference (P = 0.0265) was observed between the PBS-exposed group and the untreated group.

**Figure 6.**
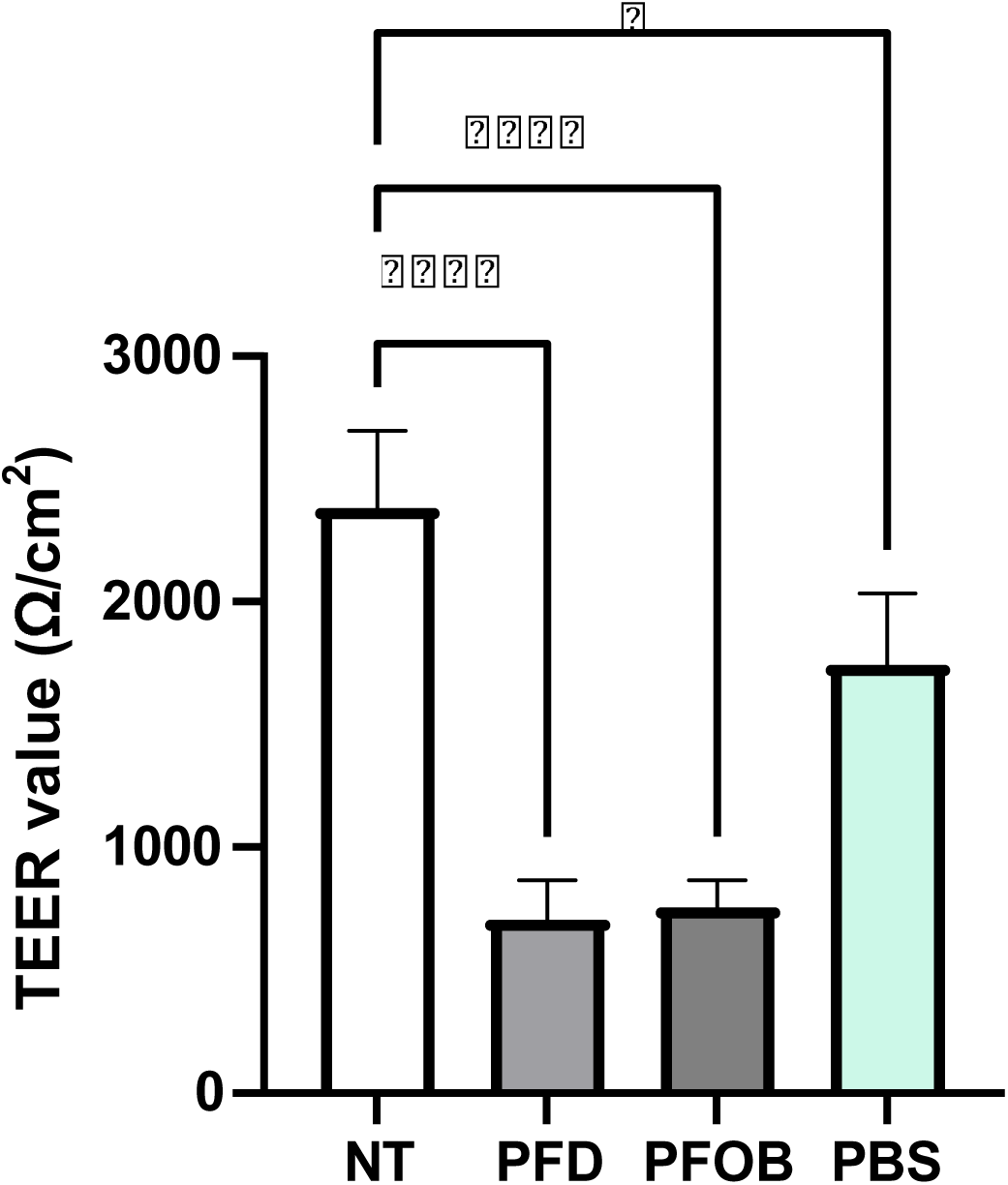
Perfluorocarbons (PFC) and phosphate-buffered saline (PBS) decrease 16HBE14o-transepithelial electrical resistance (TEER). 16HBE cells were cultured air-liquid interface in transwells. The apical compartment was filled with perfluorodecalin (PFD), perfluorooctyl bromide (PFOB), or PBS for 24 hours. The untreated group (NT) remained in the air-liquid interface. TEER values (Ω/cm2) were measured at the experiment endpoint, and data are presented as bar graphs of mean ± SEM (n = 3). One-way ANOVA was used to determine the significance of exposure conditions, with a post hoc test comparing differences between PFC and PBS groups to the untreated group. ****P<0.0001, *P<0.05.

### Assessing the effects of 24 hours PFC exposure on hSABCi cell line

#### hSABCi epithelial morphology and integrity are not deleteriously affected by PFC exposure vs the 16HBE model

Compared to 16HBE monolayer formation in transwell, hSABCi can differentiate into a bone fide pseudostratified epithelium with differentiated cell, including columnar AEC and goblet cells [37]. The visibility of hSABCi were worse (Figure 7) under brightfield microscope vs the 16HBE monolayer in the transwell system which could be attributed to a lower refractive index amenable to microscopic observations. At confluent, the hSABCi epithelial layer also possesses a similar cobblestone-like structure, though it is not homogenised or organised as compared to 16HBE in submerged culture. This may be due to these cells being immortalized in the squamous phenotype vs at terminal morphology which is a characteristic of the 16HBE model [40]. Due to the resolution of the brightfield microscope, hSABCi cells were assessed for broad epithelial layer formation, morphology, and density. Flicking motions from ciliated cells were not observable, which may reflect the low expression of ciliated cells or shorter-length cilia in differentiated hSABCi. ([37]; and in communication with collaboration – Melbourne University – see [41]).

**Figure 7.**
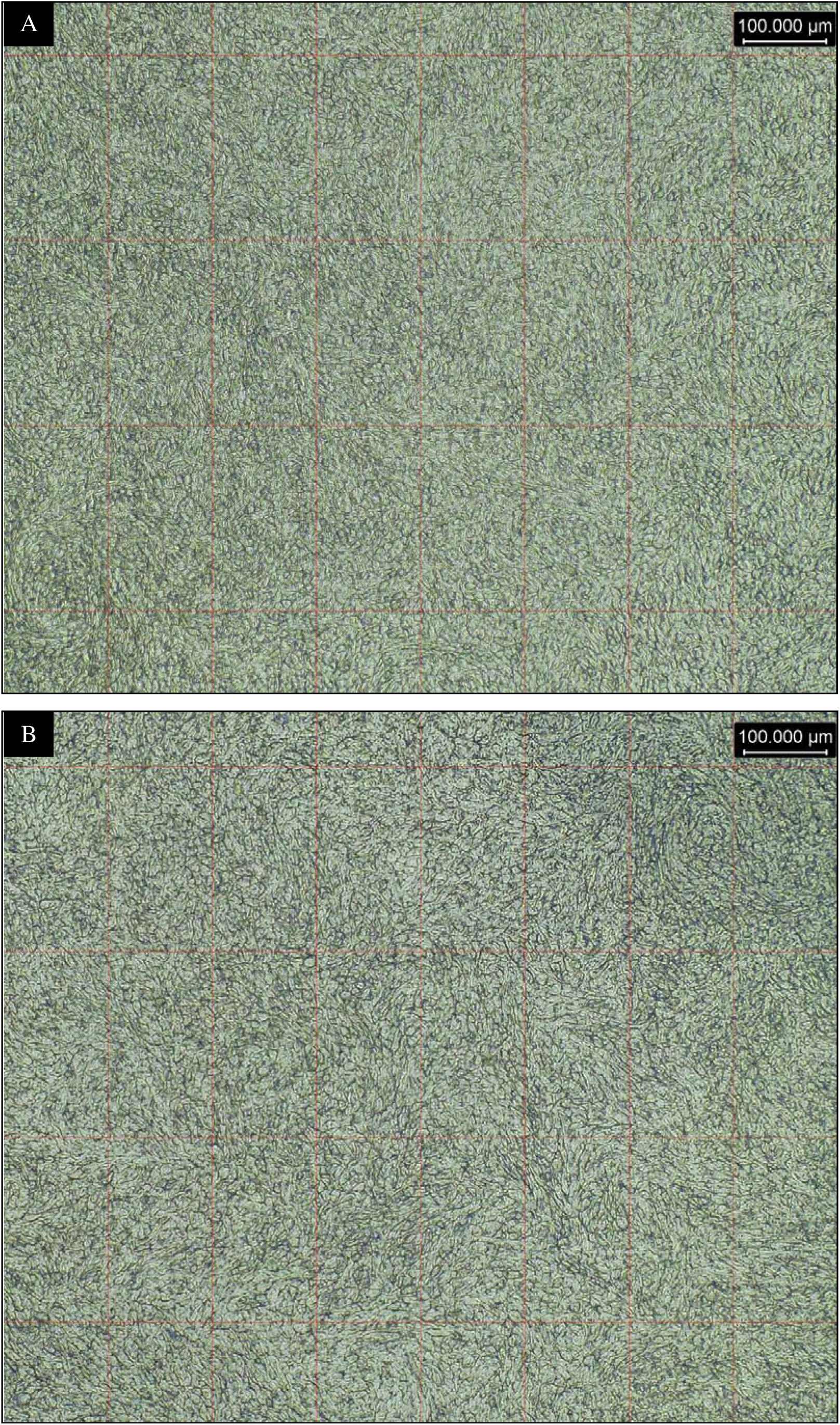
The morphology of hSABCi-NS1.1 under brightfield microscope. Pictures were taken at 10x objective lens after 24 hours of exposure experiment. Increasing magnifications decrease the visibility of cell-cell connections and were not included. (D) Air-exposed hSABCi (untreated) (E) Perfluorooctyl bromide-exposed hSABCi

Comparative microscopic examination of hSABCi before and after 24 hours of PFOB exposure showed no distinct morphological difference. Correspondingly, comparing PFOB-exposed hSABCi to untreated group revealed similar results, hSABCi maintained its heterogenous cobblestone structure with no apparent disruption in epithelial integrity and confluence. Additionally, no cell detachment, rounding, or related morphological signs of cellular stress or death were found. These findings suggests that short term exposure of PFOB has no visible effects on the structural integrity of differentiated hSABCi cells.

#### PFOB-exposed hSABCi exhibit low-level apoptotic changes and minimal modulation vs basal autophagy

PARP cleavage and autophagy markers were again used to exhibit cellular health/stress. Figure 8A shows low levels PARP cleavage in the PFOB-exposed groups, where the signal strength was almost indistinguishable from the background levels of apoptosis. Comparing levels of PARP cleavage between untreated and PFOB-exposed groups also shows no distinct differences, evident in low to no apoptotic events after PFOB exposure. This is further supported by quantitative analysis (t-test) of cleaved PARP blots, no significant differences (P = 0.6185) were found.

**Figure 8.**
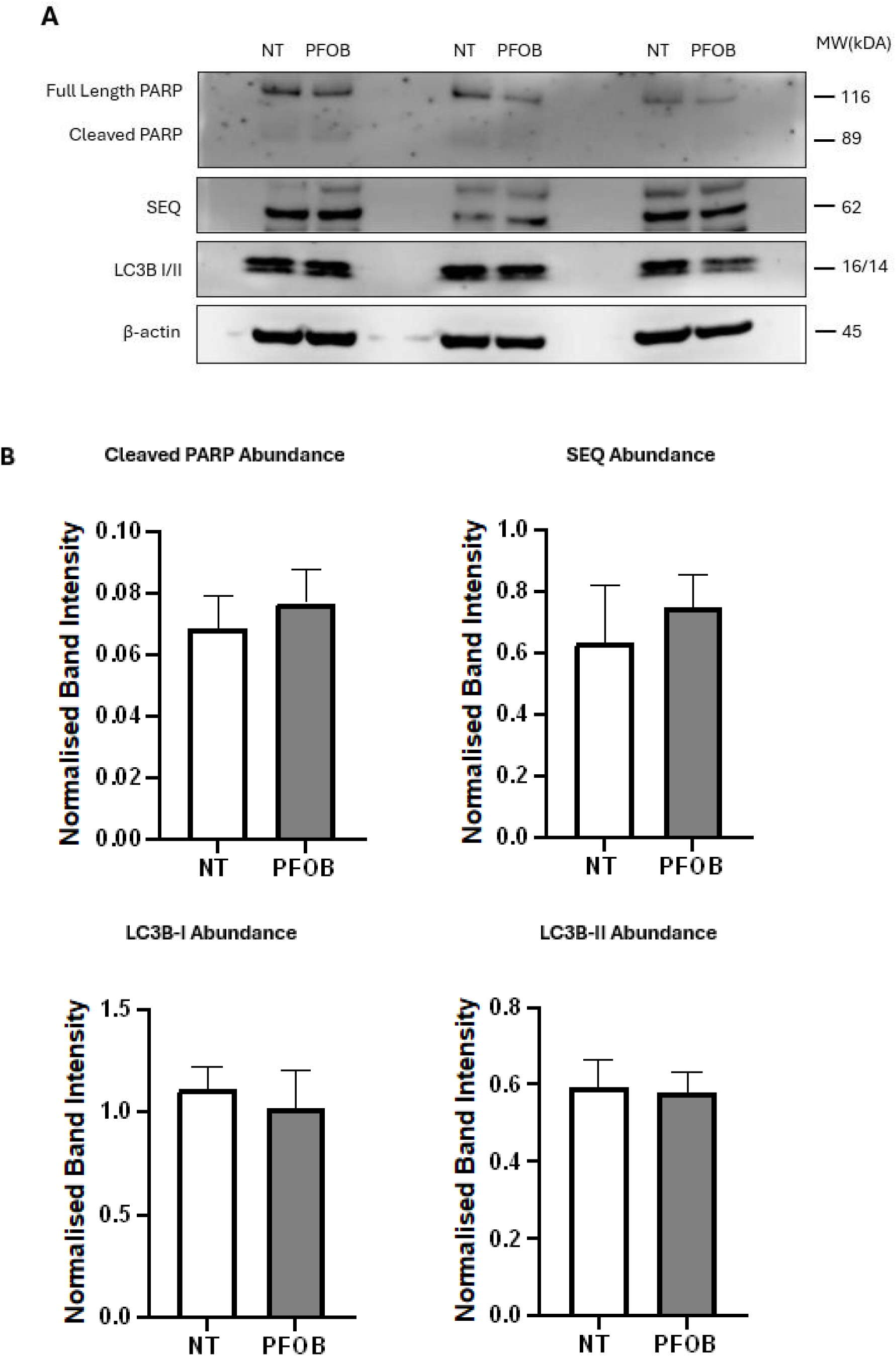
Perfluorooctyl bromide inertness results in no induction of apoptosis or autophagy in hSABCi-NS1.1 cells. (A) Western blot analysis of hSABCi-NS1.1 cells exposed to perfluorooctyl bromide (PFOB) for 24 hours. Blots were probed for full-length and cleaved PARP (apoptosis marker), SQSTM1 (SEQ), and LC3B-I/II (both autophagy markers). β-actin served as a loading control. Cleaved PARP was nearly indistinguishable from the background, indicating minimal apoptosis. Multiple bands were observed in the SEQ blot, likely representing post-translational modifications. The band with the strongest intensity was quantified. (B) Quantification of blot intensities (A) normalised to β-actin. Data are presented as a bar graph and mean ± SEM (n=3). There were no significant differences (P>0.05) between blot intensity across all biochemical analysis for untreated (NT) and PFOB group. The variance shown of autophagy markers could be attributed to biological variance, evident by overlapping error bars.

Similarly, blot intensities for signature markers of autophagy: SEQ, LC3BI and LCBII were uniform from blot observation (Figure 8A), as well as quantitative analysis showed no significant differences (P >0.05) across each of these autophagy markers (n = 3). These findings evidence the inert biological effect for PFOB and suggest this PFC has significant promise for extended investigation in the context of laboratory modelling of the airways (and relate airway diseases).

#### hSABCi treated with PFOB exhibit outcomes consistent with the air exposure barrier integrity

In addition to the microscopic examination of hSABCi overall morphology after PFOB exposure, hSABCi were further determined for outcomes related to barrier integrity. TEER measures and molecular analysis of tight junction protein abundance (Occludin-1 and claudin-1) were used as outcome measures that relate to the presence (or absence) of functional tight junction complexes Figure 9 shows a comparison of TEER values at Day 0 and Day 1 between the untreated and PFOB exposure conditions. Both groups showed a modest reduction in TEER values on day 1, and statistical analysis showed no significance in the decrease; TEER values of the untreated group were also not significantly different from those of the PFOB-exposed group (P>0.05).

**Figure 9.**
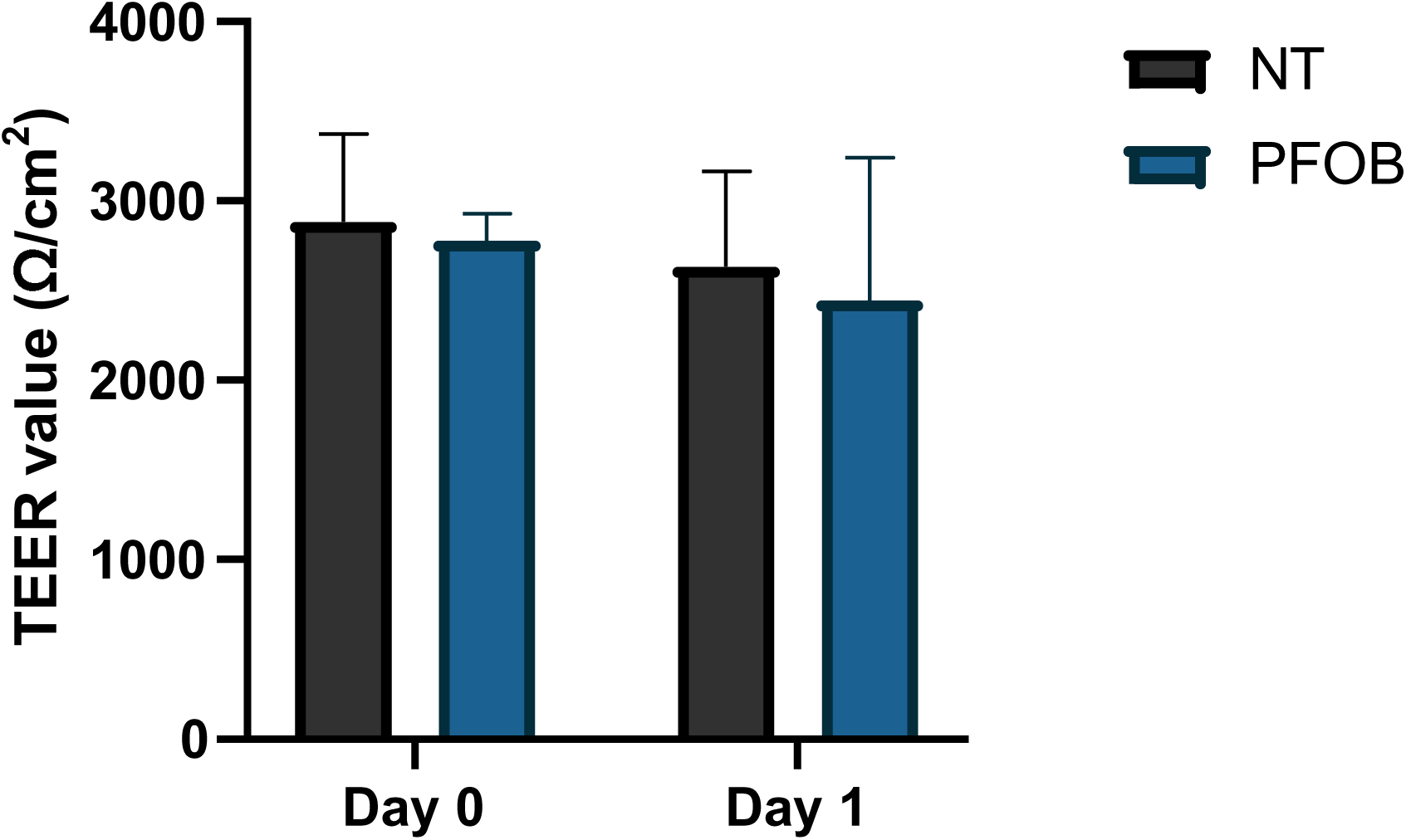
Perfluorooctyl bromide has not observable consequence for hSABCi barrier integrity. Small airway epithelial cells hSABCi were cultured and differentiated in ALI. Transepithelial electrical resistance (TEER) values (Ω/cm2) were measured before perfluorooctyl bromide (PFOB) exposure at Day 0 and the 24-hour exposure endpoint (Day 1). Differentiated hSABCi remained in the air-liquid interface and was assigned to the untreated (NT) group. Data are represented as bar graphs indicating mean ± SD. The decrease of TEER value observed from untreated and PFOB-exposed groups was nonsignificant; no significant differences were found between the untreated and PFOB-exposed groups at either timepoints (P>0.05).

Furthermore, Western blot outcomes for tight junction proteins (Figure 10) showed similar blot sizes for Occludin-1 and claudin-1 when comparing the untreated and PFOB-exposed groups. Quantifying the blot signal magnitudes showed similar intensity values that were not significantly different between sample groups for either of the tight junction proteins examined. These results are convincing evidence that indicate differentiated hSABCi cells maintains barrier integrity to a relatively short-term (24 hour) PFC exposure, as evidenced by consistent TEER values and tight junction protein expression vs the control exposures.

**Figure 10.**
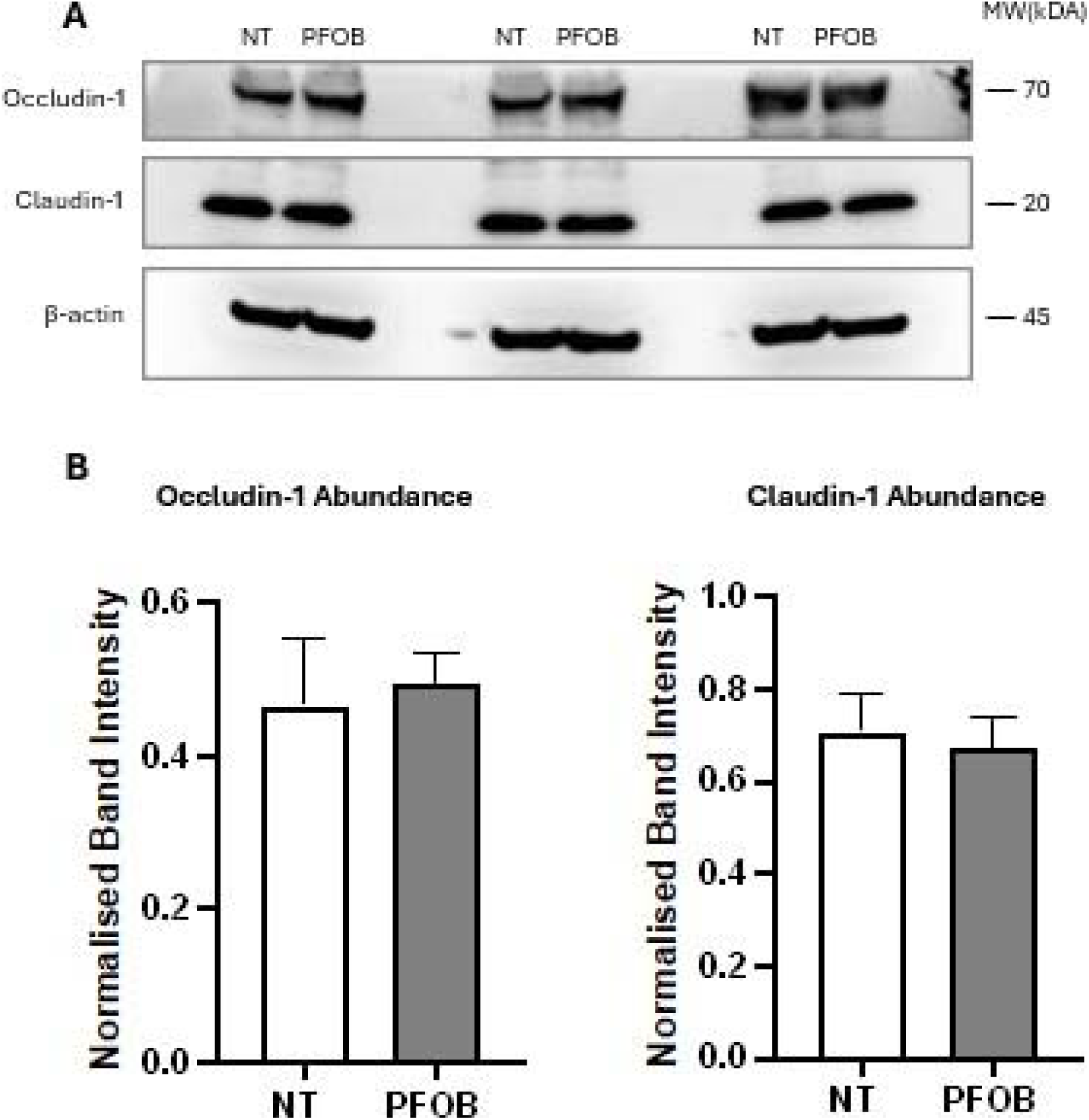
Differentiated hSABCI-NS1.1 cells shows no significant alteration of tight junction protein expression after perfluorooctyl bromide exposure. (A) Western blot analysis of hSABCi-NS1.1 cells exposed to perfluorooctyl bromide for 24 hours. Blots were probed against tight junction protein Occludin-1 and claudin-1 with β-actin serving as a loading control. (B) Quantification of blot intensity from (A). Data are presented as a bar graph representing mean ± SEM (n=3). Both tight junction protein expressions were not significantly different (P>0.05) between untreated and perfluorooctyl bromide-exposed hSABCi-NS1.1 cells. Evaluating hSABCi cellular outcomes after extended PFC exposure (72 hours) Considering the duration of PFC liquid ventilation can last up to days, we conducted PFC exposure for an extended 72 hours and assessed as per section 3.3

#### Extended PFC exposure produced no discernible changes in hSABCi cellular morphology

The microscopic examination in hSABCi exposed to PFC for 72 hours resembled observations of the 3.3 experiment module. Even after 3 days, hSABCi remained viable and retained its cobblestone structure morphology, similar to what was observed in **3.3.1** PFOB exposure (Figure7B). No signs of cell rounding, blebbing or detachment that disrupts the hSABCi epithelial integrity were observed.

#### The effect of extending PFC exposure on apoptosis and autophagy outcomes

Figure 11 Western blot analysis did not produce outcomes of visible cleaved PARP protein, and therefore quantitative analysis was not possible. This is consistent with the absence of morphological changes associated with apoptosis, which supports a conclusion that PFC exposure, even extended to 3 days, does not induce apoptosis. Moreover, Western blot of autophagy markers shows similarities between untreated and PFC-exposed groups where protein band sizes and intensities are identical. Quantitative analysis of SEQ, LC3BI and LC3BII shows that all autophagy markers have lowered band intensities in the PFC-exposed group compared to untreated control. Still, the differences are not significant, nor is it of magnitude. These outcomes suggest that hSABCi cells were not affected by the presence of apical PFC liquid, as PFC exposure did not induce apoptosis or autophagy disruption.

**Figure 11.**
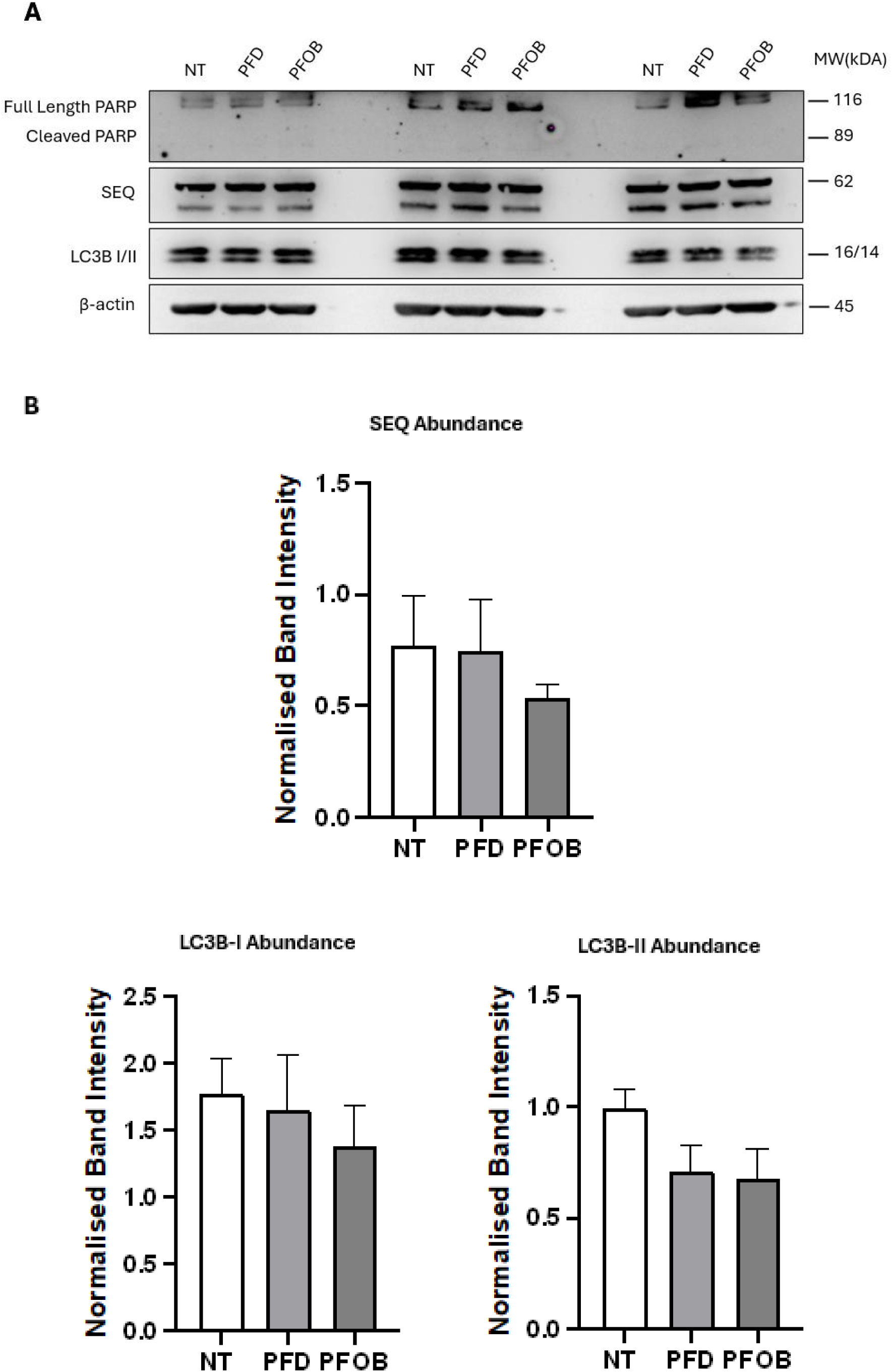
PFC inertness is reflected in hSABCi low PARP cleavage activity and basal autophagy level. (A) Western Blot analysis of hSABCi cells exposed to perfluorodecalin (PFD) and perfluorooctyl bromide (PFOB) and remained in the air-liquid interface (NT). The membrane was probed against PARP protein (full length and cleaved length), indicating apoptosis, SQSTM1 (SEQ), and LC3BI/II, both autophagy markers. No visible PARP cleavage can be observed, and all autophagy markers are identical in blot intensity between untreated and PFC-exposed groups. (B) Quantitative analysis of western blot (A). Data represents mean ± SEM (n = 3). No visible cleaved PARP protein was observed from (A) and thus was not included in the quantitative data. Untreated overall showed higher blot intensities in all autophagy markers compared to the PFC-exposed group. Further statistical analysis revealed differences were not significant (P>0.05).

#### Sustained barrier activity was observed in hSABCi cells exposed for a prolonged period to PFC

The barrier function of hSABCi was evaluated after 72 hours of PFC exposure by measuring TEER. The TEER values were recorded at the experiments’ endpoint, and comparative assessments were made between the untreated group (the air-only condition) and the PFC-exposed groups. Figure 12 shows a slight increase of TEER values in PFC-exposed cells, though the increases observed were not significantly different (P<0.05).

**Figure 12.**
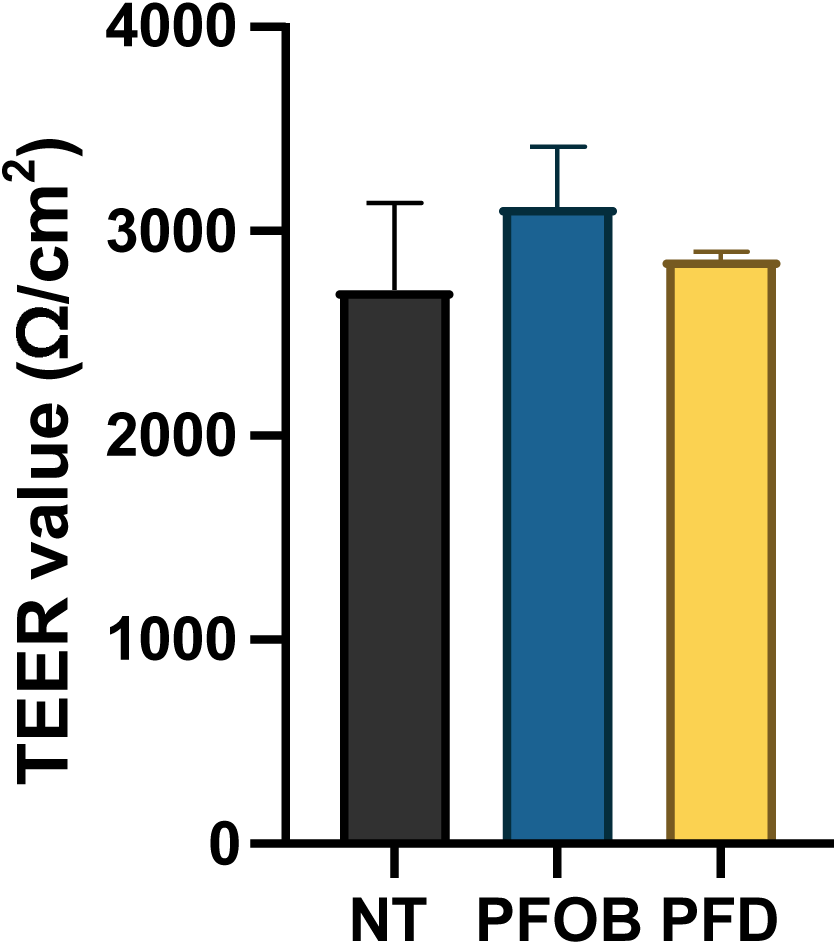
Small airway epithelial TEER maintenance after prolonged PFC exposure. Before PFC exposure, small airway epithelial cells hSABCi were cultured at an air-liquid interface. The cell line maintained barrier integrity as indicated by TEER measurement at the end of 72 hours of exposure. Both perfluorooctyl bromide (PFOB) and perfluorodecalin (PFD) maintained TEER levels comparable to the untreated group at both time points. Data are presented as bar graphs reflecting mean ± SD (n = 3). No statistically significant differences were observed between any groups.

To further assess the barrier function, we performed a western blot to analyse tight junction protein expressions. Figure 13A demonstrates a significant increase in ZO-1 protein expressions in PFC-exposed groups compared to untreated hSABCi. Quantitative analysis of blot intensities revealed a pattern of increased ZO-1 in PFOB and PFD groups. One-way ANOVA results indicate that the exposure condition is significant (P = 0.0175) in increasing ZO-1 protein expression, with the PFD group showing a particularly significant increase (P = 0.0114) relative to the untreated group, with an almost 2-fold increase in blot abundance. In contrast, Occludin-1 and Claudin-1 expressions were not significantly different in the PFC-exposed group compared to the untreated group, and the observed variations were within the range of normal biological variations.

**Figure 13.**
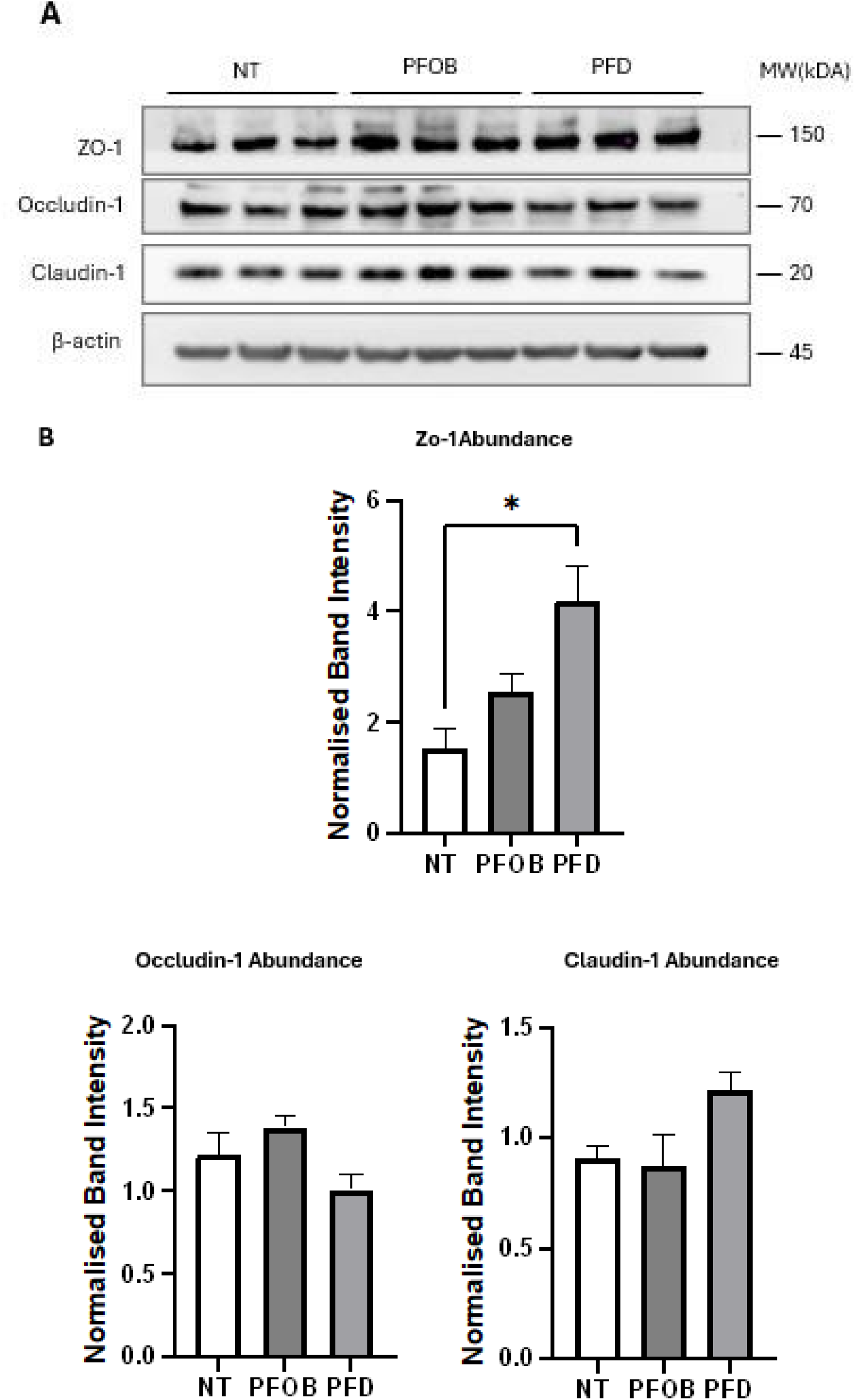
Stable tight junction protein expression in hSABCi after prolonged PFC exposure. (A) The expression of three tight junction proteins, ZO-1, Occludin-1 and Claudin-1, was assessed using western blot. hSABCi cells were exposed to perfluorooctyl bromide (PFOB), perfluorodecalin (PFD), or air (NT). ZO-1 expression appeared to be upregulated in the PFC-exposed groups, while Occludin-1 and Claudin-1 blot intensities are similar across all groups. (B) Quantification of tight junction protein western blot (A). Blot intensities were presented as mean ± SEM (n = 3). One-way ANOVA analysis revealed PFC exposure had a significant effect (P = 0.0175) in ZO-1 blot intensities, and post-hoc indicated only the PFD group resulted in significant increases (P = 0.0114) in ZO-1 expression compared to the untreated group. No other significance was found in Occludin-1 or Claudin-1 (A) quantification. ZO-1 blot results show a pattern of increased expressions in PFC-exposed groups, while the others showed a more uniform expression.

### Assessing the effects of prolonged PFOB & PBS exposure on primary human nAEC

#### Primary nasal AEC (nAEC) epithelia is preserved after exposure to PFOB

Primary nAEC cells exhibit epithelial cobblestone structure as seen in 16HBE and differentiated hSABCi. However, nAEC also exhibit full mucociliary differentiation; this phenotype is organotypic and distinct to the immortalized cell lines that were previously assessed. Mucus production was apparent, and as there is no expectoration (as per in mammals) of the accumulated mucins, these cultures require PBS clearance/washing to clear excess mucus build-up. Cilia beating kinetics were readily evident and visualised with routine brightfield microscopy, especially in areas where undulations of the topology of the epithelial layer(s). Undulations refer to the “hills and valleys-like” topography of nAEC culture, which was more pronounced in nAEC cells and necessitated readjustments of the focal point to maintain clear cellular observations. This contrasted with the uniform monolayer of the16HBE model, where a single focal plane is was generally sufficient for observing the entire cell layer. This speaks to the variable nature of primary samples and the formation of a three-dimensional epithelial morphology with basal cell supporting the differentiation of various specialised cell types, such as columnar, goblet and tuft cells [1].

While performing routine microscopy for the nAEC model during the exposure interval, cilia beating remained readily evident across all groups. The PBS control exposure (maintained in parallel to the PFC exposures as a control for “a non-air liquid”) presumably was able to dissolve mucus effectively. This was evidenced by straightforward identification of cilia activity for this condition. Furthermore, the cobblestone structure was maintained in all groups, resembling hSABCi cells, where the cobblestone structure is less organised than that of 16HBE cells. Interestingly, the accumulation of mucus production did not result in difficulty visualising nAEC. PBS-exposed nAEC exhibited signs of cell-to-cell dissociation and morphological change, visible at 40x magnification, and lower magnifications showed dissolved mucus that appeared as black streaks. Irrespective, each condition for the nAEC model maintain structure integrity apart from cell rounding on day 3. No apparent disruption of the epithelial layer, such as gaps in cell growth observed in 3.1.1 16HBE cells, was found.

#### Baseline PARP protein and autophagy modulation observed in PFOB-exposed primary nAEC

Western blot results from Figure 14A show similar PARP cleavage between nAEC cells in air (untreated) and PFOB exposure; no cleaved PARP nor full-length PARP can be observed in the exposed nAEC group. This may be due to the apoptotic signal being accumulated in detached cells, and the methodology of aspirating and refilling apical fluid every day during the experiment resulted in accumulated apoptosis signal loss. Nevertheless, PFOB exposure resulted in outcomes similar to previous modules (see **3.4.2**) where the cleaved PARP signal was comparable (P>0.05) to the basal level in the untreated group. Conversely, SQSTM 1 is highly upregulated (P<0.0001) compared to untreated control and may indicate upregulated autophagy; this is supported by the conversion of LC3BI to LC3BII. The variations in LCBI/II in the exposed group were comparable to untreated control, and combined with low expression of SQSTM1, autophagy is highly unlikely to be modulated as a result of prolonged exposure to PFOB.

**Figure 14.**
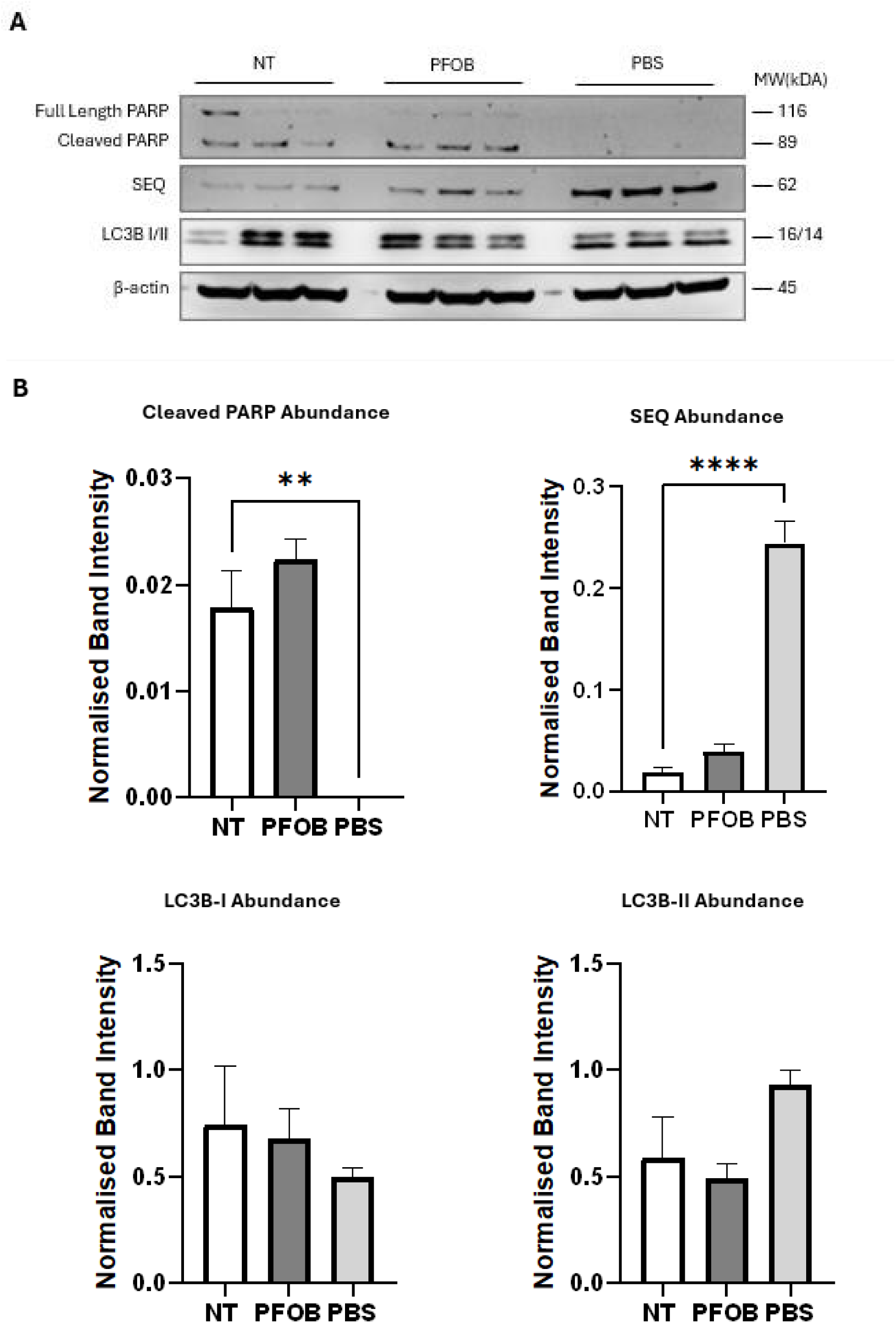
Western blot analysis reveals absence of PARP protein and upregulation of SQSTM 1 in PBS-exposed nAEC cells. (A) Western blot outcomes for Primary nAEC cultured in ALI condition and exposed apically to PFOB and PBS. The membrane was probed for PARP (apoptosis marker), SQSTM1 (SEQ) and LC3BI/II (both autophagy markers). Cleaved PARP protein intensity is comparable between untreated and PFOB-exposed nAEC. However, lanes assigned for the PBS-exposed sample showed no PARP protein bands except for some background dots. In contrast, SQSTM1 blots in the PBS group showed the most substantial blot intensities, followed by the PFOB group and the lowest was observed in the untreated group. Moreover, LC3BI shows an overall decrease from untreated to PFOB group, then PBS group in order; LC3BII blot is the highest intensity in the PBS-exposed group, while untreated and PFOB group are similar. (B) Quantification of apoptosis and autophagy protein blots (A). Data are presented as mean ± SEM (n = 3). The absence of PARP protein resulted in a significant difference (P = 0.0025) in cleaved PARP protein between the PBS and untreated groups. In contrast, the PFOB group showed comparable results that were not significant. Furthermore, SQSTM 1 protein is significantly (P <0.0001) more expressed in the PBS group than in the PFOB or untreated group. No significance was found in LC3BI/II blots, and no significant differences were identified between all groups.

#### Primary nAEC barrier function is preserved after extended PFOB exposure

TEER of nAEC cells were taken over the 72-hour exposure period and compared between untreated, PFOB and PBS-exposed groups. As shown in Figure 15, untreated nAEC cells display TEER values similar to PFOB-exposed, with no significant differences observed in this comparison. In contrast, PBS deleteriously affected nAEC barrier integrity rapidly, as shown in Figure x, where the TEER values decline from day 1 and plateau over days 2-3. Day 1-3 TEER values from the PBS-exposed group were significantly lower than the respective untreated group, with day 2 being the most significantly different (P values range from <0.05 to <0.001, vs the air/control exposure).

**Figure 15.**
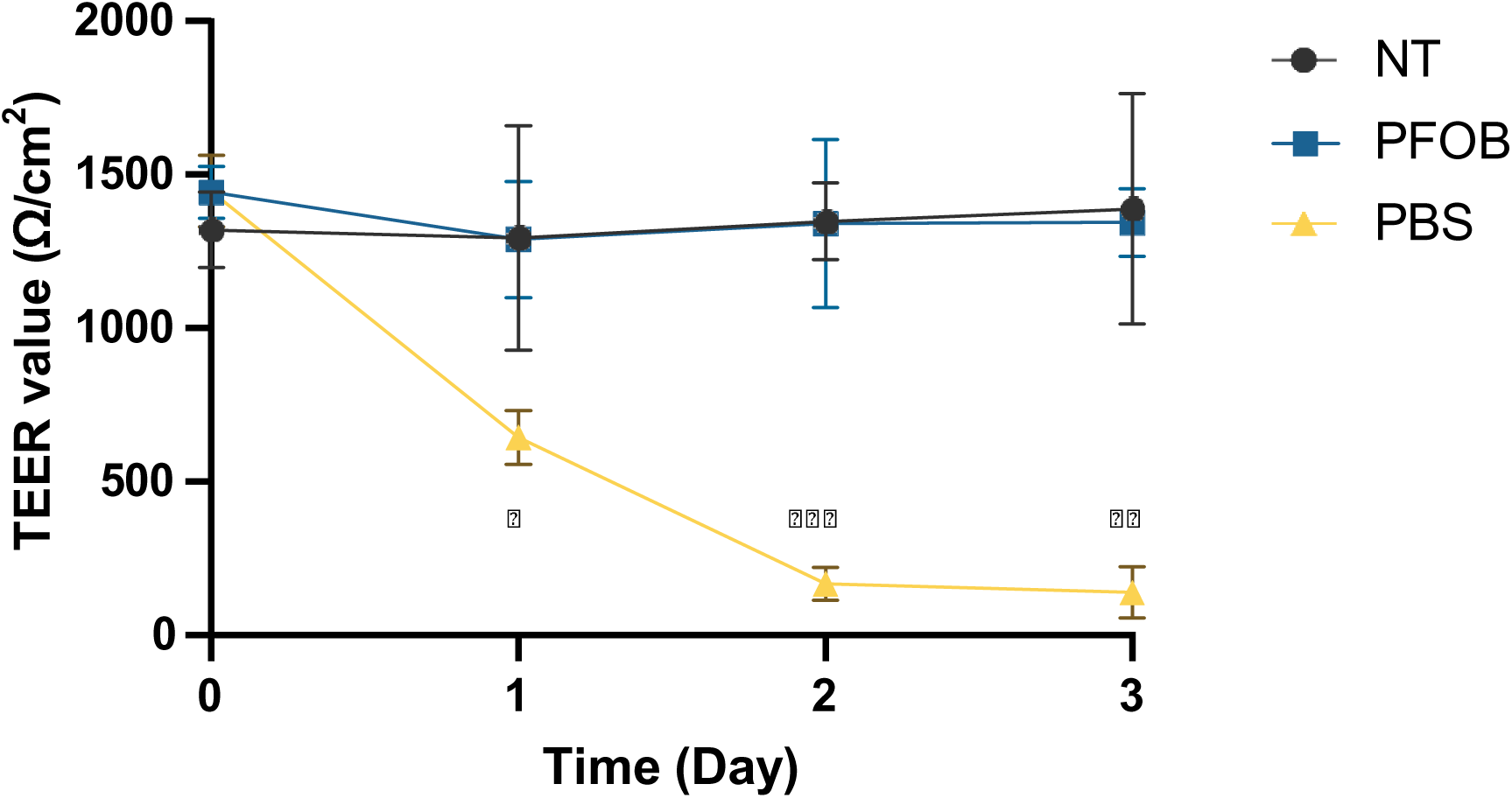
Primary human nAEC maintains barrier integrity following perfluorooctyl bromide exposure but not PBS. The electrical impedance of differentiated nAEC cultured at an air-liquid interface was measured during 3-day exposure to perfluorooctyl bromide (PFOB) and phosphate-buffered saline (PBS). PFOB-exposed nAEC maintained TEER levels comparable to the no-treatment (NT) control, while PBS significantly reduced the barrier integrity of nAEC. Data are presented as mean; error bars represent 95% confidence intervals (CI) from 3 independent experiments. Statistical significance was determined by two-way ANOVA followed by Dunnett’s multiple comparisons test. The Dunnett’s test confirmed that PBS significantly reduced TEER compared to NT at all time points, while no significant differences were observed between PFOB and NT. Significant differences are indicated as *p < 0.05, **p < 0.01, ***p < 0.001.

This disruption of barrier function is further supported (Figure 15) by tight junction protein Occludin-1 and Claudin-1 expression from western blot analysis. Occludin-1 expression was upregulated in PBS, and Claudin-1 protein was not detectable in the PBS group. The Occludin-1 increased expression is significant (P<0.05), and the Claudin-1 quantification analysis was impossible. In the other exposure group, Occludin-1 expression was even more significant (P = 0.0065) in PFOB exposure than in PBS, but Claudin-1 blot intensities in this group did not significantly differ from untreated control. These findings suggest that PBS severely compromise nAEC barrier function, as evidenced by a significant decrease in TEER values and dissolution of tight junction (Claudin-1 absence). One explanation for this is that PBS (vs the biologically inert PFC conditions) actively dissolves and therefore removes the protective effect imparted by the airway surface liquid and mucin layers that are central to primary epithelial cell integrity and cilia beat activity. Nonetheless, PFOB biocompatibility is once again demonstrated through preserved nAEC barrier function after extended exposure.

#### Primary nAEC IL-6 and IL-8 secretion in transwell after PFOB and PBS exposure

Cytokines IL-6 and IL-8 secretions were assessed by performing ELISA on apical wash and basal media samples of nAEC that were either untreated or exposed to PFOB or PBS.

#### IL-6 Secretion

Figure 17A shows that PFOB exposure did not induce significant IL-6 release compared to the untreated control in either apical wash or basal media. PBS exposure, on the other hand, resulted in a marked increase in IL-6 levels in the basal media, although this difference was not statistically significant compared to the untreated control.

**Figure 16.**
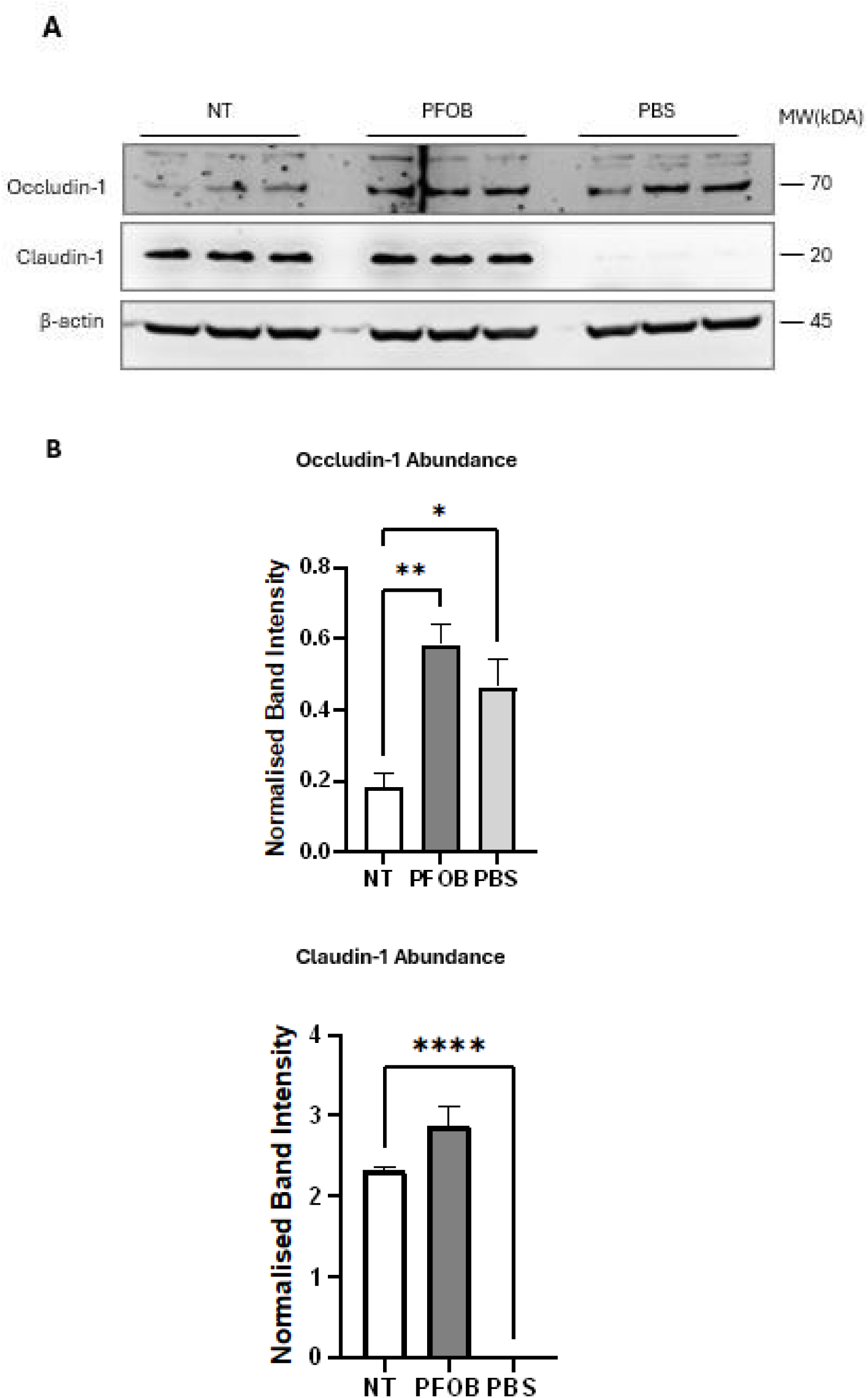
Primary nasal airway epithelial cells significantly altered tight junction protein expressions with apical liquid interface. (A) Western blot analysis of tight junction protein markers in primary nasal airway epithelial (nAEC) cells after 72 hours of liquid exposure in transwells. Blots were probed for Occludin-1 and Claudin-1 with β-actin as a loading control. Exposure conditions were untreated (NT), perfluorooctyl bromide (PFOB) and phosphate-buffered saline (PBS). (B) Quantification of tight junction protein blots intensity. Data are presented as mean ± SEM in a bar graph format. PFOB exposure significantly increased occludin-1 expression (P<0.01) but not claudin-1, compared to the untreated group. Conversely, PBS exposure also significantly increased Occludin-1 expression (P<0.05) but substantially decreased (P<0.0001) Claudin-1 expression compared to the untreated group. As the Claudin-1 blot was not detectable in (A) for the PBS group, it is represented as 0.

**Figure 17.**
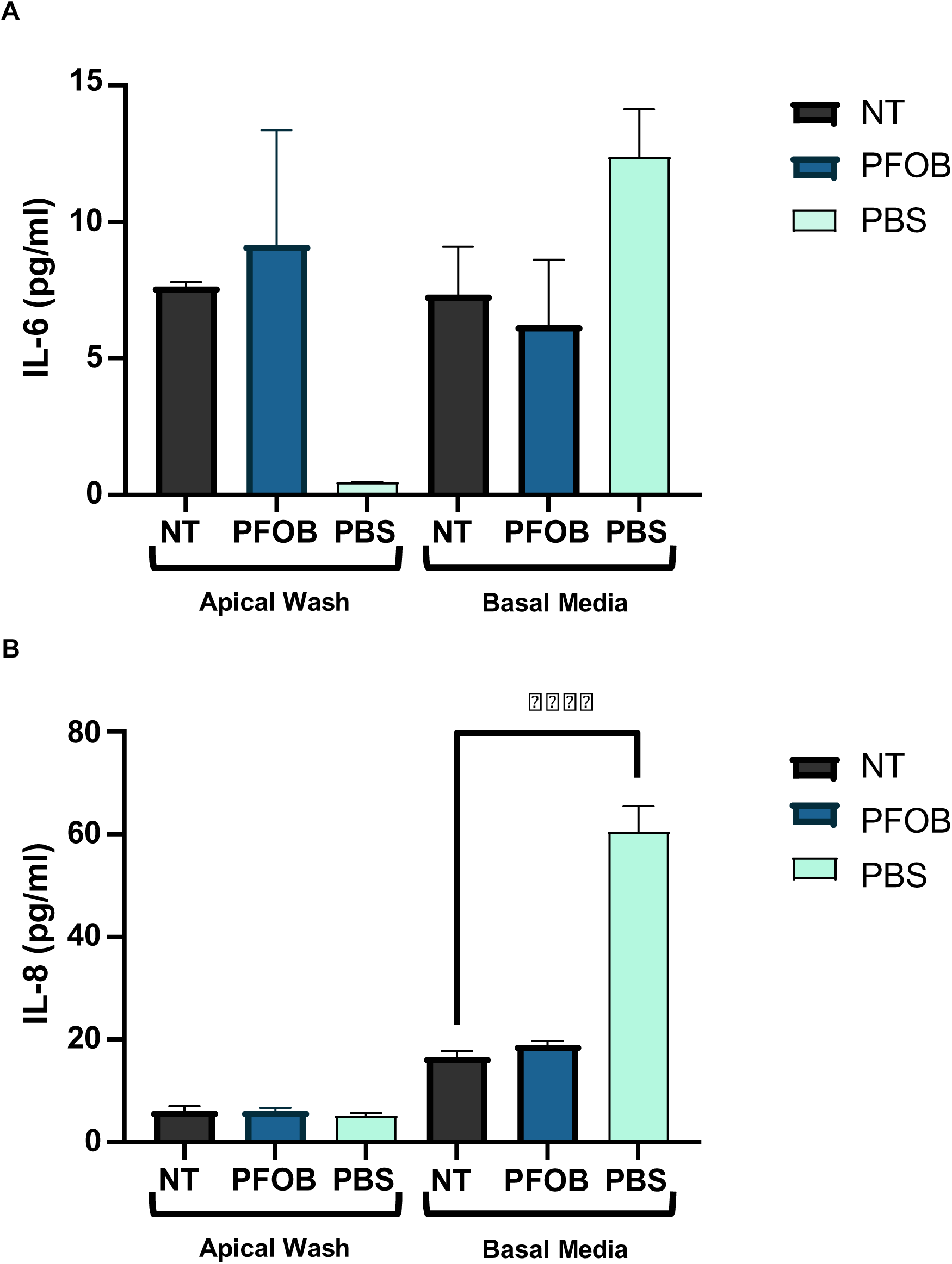
Primary nAEC IL-6 and IL-8 levels after PFOB and PBS exposure. Differentiated nAECs were exposed to perfluorooctyl bromide (PFOB) and phosphate-buffered saline (PBS) for 72 hours. ELISA assessed IL-6 and IL-8 secretion in apical washes and basal media. Data are presented as mean ± SEM represented by bar graphs. (A) IL-6 levels between untreated and PFOB are comparable from apical wash or basal media. The PBS group shows low levels of IL-6 apically but substantially more IL-6 in basal media. Despite that, the differences between exposure groups (PFOB & PBS) and untreated reference were nonsignificant. (B) IL-8 levels appear uniform across all apical wash samples, and corresponding levels in basal media are all higher. While untreated control and PFOB show comparable IL-8 levels, IL-8 detected in the PBS group’s basal media is considerably increased, showing high significant differences (P<0.0001) in the untreated group.

#### Il-8 Secretion

Apical IL-8 levels (Figure 17B) were uniformly low across NT, PFOB, and PBS groups. However, ELISA of basal media reveals elevated IL-8 levels across all conditions. Most notably, the PBS group exhibited substantially more IL-8 secretion, roughly 3 times compared to the other groups, and statistical analysis revealed (P<0.0005) differences when compared to the untreated group.

These data suggest that PFOB inertness is further displayed through similar cytokines secretion between the nAEC untreated control and the PFOB-exposed group. In contrast, PBS exposure resulted in pronounced IL-8 secretion, which suggests that PBS caused some levels of cellular stress in nAEC.

## Discussion

### Summary

This thesis investigates the effects of perfluorocarbon (PFC) exposure on human airway epithelial cells (AECs) in vitro. Firstly, the necessity of a complex 3D model for the PFC exposure experiment was determined by comparing apoptosis outcomes in a conventional submerged system and an organotypic air-liquid interface (ALI) system in 16HBE14o- (hereafter referred to as 16HBE) cells. Immiscibility between media and PFC presents a limitation of the conventional submerged system. Hence, ALI architecture was employed for subsequent PFC exposure experiments. The architecture of the ALI system overcomes the issues of providing nutrients to the cells while exposing to the PFC. As a result, near absence cleaved PARP (as a readout for apoptosis) was observed across 16HBE, hSABCi-NS1.1, and primary nasal AEC exposed to PFC at an ALI, which indicates limited apoptosis-inducing effects within the exposure period (24 and 72 hours).

Similarly, autophagy (a sensitive readout for cellular homeostasis and nutrient availability) remained at a basal level across the n=3 airway culture groups, evidenced with comparable SQSTM1 and LC3BII abundance between untreated AECs and the PFC-exposed group. Moreover, PFC exposure decreased transepithelial electrical resistance (TEER) measurement (a reliable proxy measure for whole epithelium integrity) in 16HBE cells but not in hSABCi and nAEC culture, even after prolonged (72 hours) PFC exposure. The preservation of barrier activity was also reflected in tight junction protein abundance (Occludin-1 and Claudin-1) shown in Western Blot. This discussion section will interpret experimental findings in the context of current literature, list limitations in the study and suggest future research directions

### Experimental results interpretation

#### Cleaved PARP-1 as signatures of apoptosis

Programmed cell death, or apoptosis, is a complex and highly regulated process that removes unwanted or damaged cells to maintain tissue homeostasis [42]. It is characterised by biochemical and distinct morphological features, such as cell shrinkage, rounding, membrane blebbing, and pyknosis (chromatin condensation) [43]. Signals from within (intrinsic) cells and from the surrounding (extrinsic) environment initiate apoptosis and lead to a cascade activation of essential effector proteases (caspases) that execute cell death. In this thesis, apoptosis initiation in the form of anoikis is observed in the 16HBE submerged model. Anoikis is cell death by detachment from the extracellular matrix [44] and is common in in vitro models as there are no inflammatory cells to clear detached cells and the substratum is removed from the sub-epithelial basement membrane (to name one factor).

**Poly (ADP-ribose) polymerase**, or PARP, is a family of proteins that repair DNA damage [45]. PARP-1 is the most studied member and is identified for its central role in DNA repair. Upon detection of a single-strand break, for example, PARP protein, through activation, binds to the site of damage and facilitates the recruitment of DNA repair proteins (such as DNA ligase III) for the repair process [46]. During apoptosis, however, the 116-kDA PARP-1 protein will be cleaved by activated caspases 3 and 7, leaving behind 89-kDA fragments consisting of central (auto modification) and C terminal (catalytic) domain [47]. The inhibition of DNA repair mechanisms via PARP protein fragmentation is a well-accepted hallmark of apoptosis [47].

This study evaluated cellular damage and responses by measuring PARP-1 cleavage in human AEC following PFC exposure. Firstly, the apoptosis (anoikis) observed in the 16HBE submerged system was starvation-induced due to the model’s limitation. This was supported by a similar PARP cleavage pattern in a Western blot of intentionally starved 16HBE (Figure 3). Organotypic cultures (16HBE14o-, hSABCi-NS1.1, and primary nAEC) were employed and grown at an air-liquid interface to overcome this limitation. PFC exposure resulted in no apparent apoptotic morphological changes such as cell detachments, rounding and blebbing via routine light microscopy. PARP protein Western blot also showed either a near absence of cleaved PARP or comparable signals to untreated controls (AEC cultures exposed to air) that indicate basal PARP activity. Hence, limited PARP cleavage following PFC exposure strongly supports PFC as a biocompatible agent, as shown in liquid ventilation clinical trials and animal studies where demise mainly was attributed to technical error (e.g. pneumothorax, see [27]), administration strategy or condition severity [20, 21]. Hypothesis 1 is supported by a polarised culture that mimics the airway architecture, which enables a more objective assessment of the PFC exposure condition, in practical terms, and as this relates to a system that closely approximates the in vivo situation for these cells. The traditional submerged system is limited to studying immiscible exposure conditions with culture media essential for survival. Moreover, hypothesis 2 is supported via modest cleaved PARP abundance as AECs in the organotypic model were unaffected by PFC exposure. This is underscored by many in vitro experiments where PFC is used in systems that do not require nutrition and/or use very brief exposure periods [48].

#### Baseline autophagy in AEC after PFC exposure

Autophagy is translated into self-engulfment or self-eating in Greek. It is a highly conserved cellular process that achieves cellular homeostasis for development, differentiation and survival [49]. It should be noted that all instances of autophagy discussed in this thesis refer specifically to the canonical pathway of macroautophagy. Briefly, the autophagic process [50] is activated by:

1. upstream signalling of cellular stress (starvation and damage) and initiated by assemblance of Unc-51-like kinase 1 (ULK1) complex with autophagy-related proteins (ATGs).
2. Initiation will result in membrane nucleation and formation of early stage phagophore
3. The phagophore will continue to expand as cytoplasmic cargos (macromolecules) are guided towards it and autophagosome is formed when the expansion complete.
4. Cargo within the autophagosome will be degraded after lysosomal fusion, and degraded product will be released into the cytoplasm.

**Sequestosome-1** (abbreviated as SEQ) is one of the key markers of the autophagic process. It is an autophagy receptor that binds ubiquitinated cargos to expanding phagophore (3) [50, 51]. Another key protein marker is **microtubule-associated proteins 1A/1B light chain 3B** (LC3B) that exist in two forms: LC3BI is formed through ATG4B cleavage of pre-LC3BI protein from the C-terminus, and LC3BII that is conjugated (by ATG3 and ATG7) with lipid molecule phosphatidylethanolamine (PE) [50].

**LC3BII** is incorporated into autophagosomes during (3) and interacts with cargo receptor SEQ, which acts as an adaptor protein that links and anchors cargo to the autophagosome [52]. The upregulation of LC3BII indicates an increased number of autophagosomes, while the upregulation of SEQ indicates increased cytoplasmic cargo for degradation during times of cellular stress.

The consequence of PFC exposure in hSABCi and nAEC was that no apparent autophagy modulation was indicated by SEQ and LC3BII abundance, which resembled baseline autophagy of the untreated control group. This is strongly contrasted with PBS-exposed primary nAEC (Figure14B), where SEQ and LC3BII were significantly more abundant, and an LC3BI exhibited a conversion pattern to LC3BII from Western Blot. PFC’s inert nature is again demonstrated in autophagy assessment, such that apical liquid like PBS results in significant stress response while PFOB exposure did not affect autophagy. These autophagy results support the second hypothesis in conjunction with apoptosis results that no decrease of cellular viability was evident from close to absence of apoptotic anoikis and autophagy level that represents basal/ biological variations in

#### Human AECs preserved barrier function following PFC exposure

TEER is widely applied to quantitatively measure barrier integrity of polarised epithelial/endothelial cell culture models. Measuring TEER can be performed in real-time without deleterious effects on cell culture and is generally based on measuring ohmic resistance.

TEER results of 16HBE in transwell following PFC exposure showed high TEER reading in the untreated group with 2380 ± 315 Ω/cm2. After unit conversion, the baseline readings of 16HBE are similarly high, with TEER values observed in [53] (TEER ≈1800 Ω/cm2) and [54] (TEER ≈ 2100 Ω/cm2). In contrast, 16HBE exposed to PBS for 24 hours did show decreased TEER of 1744 ± 289 Ω/cm2; even though the difference was significant (P<0.05) compared to untread control, the TEER value is still within the range of TEER observed in the literature [53]. After the experiment, all PBS exposures (n = 3) turned slightly pink, which may suggest basal media diffusion across the cell layer as reflected in decreased TEER. In contrast, perfluorodecalin (PFD) and perfluorooctyl bromide (PFOB), the two PFC exposure groups, saw a significant decrease (P<0.0001) in TEER value to sub-1000. This suggests that PFC exposure may penetrate the integrity of the 16HBE epithelial barrier and compromise cell-to-cell adhesion. Similar observations were made in A549 cells, where the alveolar cells responded to PFC through actin cytoskeleton remodelling and weakened adhesion [33]. Interestingly, The media permeate phenomenon seen in PFC-exposed groups was not seen in PFC groups. This may be attributed to the PFC-cell interface where PFCs’ higher density limit media leakage towards the apical compartment, preventing diffusion.

Performing similar experiment using the hSABCi model (without PBS exposure) saw opposite outcomes, where TEER value remained high and stable above 2500-3000 Ω/cm2, within the range reported in [37] of various hSABCi passages. TEER differences between untreated and PFC-exposed groups at either 24 or 72 hours were linked to biological variance due to overlapping error bars during data analysis. This result contrasts sharply with decreased TEER in the 16HBE monolayer. The TEER stability suggests a pseudostratified epithelium like hSABCi that is more representative of the bronchial airway region and expresses functional surface liquid expression, may be resistant to PFC-induced barrier disruption and a more accurate representation of PFC-airway epithelium interaction.

Despite similar culture methods to [1], TEER measurement was performed without the same apical fluid, basal media and acclimatisation period before reading TEER. Hence, the baseline TEER of primary nAEC exposed to air (untreated control) reached 1300 Ω/cm2 (Figure 15). Regardless, PFOB exposure did not disrupt barrier function, as evidenced by identical TEER values to untreated control over the 72-hour experiment period. In contrast, PFC was used as a negative control in the module due to barrier disruption in 16HBE; the decrease in TEER was substantial since day 1 and plateaued out to a low value at the end of the experiment. This further supports hypothesis 1 on the necessity of an organotypic model that allows for 3D differentiation architecture of the airway (nasal or bronchial) when assessing PFC effects on airway epithelium and hypothesis 2 of limited barrier disruption from PFC exposure.

Moreover, Western blotting tight junction protein expressions saw a similar pattern where Occludin-1 and Claudin-1 abundance were uniform between untreated groups and PFC-exposed groups of hSABCi and nAEC (Figure 13 and 16). Two outliers of significantly increased Occludin-1 and ZO-1 abundance were observed in nAEC and hSABCi, respectively, following PFC exposure (P<0.05). There was a discrepancy between the increase of tight junction protein and the lack of TEER value fluctuations. This suggests that hSABCi and nAEC may have region-specific mechanisms for regulating tight junctions as a response to PFC exposure that is not reflected in the ionic permeability of the airway epithelium. Nevertheless, barrier function and integrity were persevered after PFC exposure and support hypothesis 2

#### IL-6 and IL-8 secretions

Interleukin-6 (IL-6) and interleukin-8 (IL-8) are important cytokines involved in immune response and inflammation within the lung environment. In brief, IL-6 stimulates inflammatory cytokine production, promotes T helper cell differentiation, and contributes to tissue repair and regeneration [55]. Elevated IL-6 contributes to airway hyperactivity and remodelling, contributing to disease pathogenesis in asthma and chronic pulmonary obstructive disease [56]. IL-8 on other hand is a primarily a chemokine that recruits immune cells such as neutrophils to the site of damage or inflammation to elicit a stronger immune response against a pathogen or to assist wound healing [57]. Significantly elevated IL-8 levels are prognostic biomarkers for lung disease severity, such as the hyperinflammation in acute RDS [57].

The primary nAEC was the subject of investigation for cytokine release, with ELISA tests conducted on apical washes and basal media. The use of PBS as a positive control for barrier disruption was reflected in the increased levels of IL-6 and IL-8 from the basal media. The outcomes of the IL-6 and IL-8 ELISA tests followed a similar pattern to the previous assessments of cellular viability and barrier function, with no significant differences (P>0.05) found between the cytokine levels in the untreated control and nAEC after PFOB exposure. This suggests that PFOB does not alter the secretion of IL-6 and IL-8 in healthy nAEC, which may have implications for the use of liquid ventilation outside of medical applications for injured or diseased lungs.

### Limitations

#### Technical limitation – PBS apoptotic outcomes in primary nAEC

Firstly, PBS exposure was used as a control where negative cellular viability and barrier function outcomes were expected. Western Blot outcomes of PARP protein reveal the absence of PARP protein, which doesn’t make sense considering cell rounding and unhealthy morphological observations from routine microscopy. As the cell death measured in this experiment was of anoikis, where apoptosis occurs after cell detachment from the extracellular matrix, aspirating and replacing PBS exposure daily may have led to a loss of apoptotic signals. This was reflected in protein BCA assay results, where there was a clear loss of protein concentration. Either leaving the PBS exposure untouched during the exposure period or collecting PBS after each replacement may aid in limiting signal loss.

#### Limited exposure condition

This study did not fully explore the potential influence of ventilation parameters. During liquid ventilation, oxygenated PFC flows in the patient’s lungs with positive pressure ventilation irrespective of the type of liquid ventilation (total or partial). The PFC used in this thesis was not highly oxygenated, and the model employed was unable to allow the dynamic flow of apical liquid. Combined with mechanical stress, the interactions between PFC and airway epithelium may differ according to the condition (static at 37°C and 5% CO2 atmosphere) tested. Introducing highly oxygenated PFC exposure would also mean checking mitochondria’s function and determining whether cells undergo oxidative stress.

Secondly, while the 24-72 hours exposure period is relevant to liquid ventilation duration in clinical trials, this is still a short-term examination of PFC effects, and the long-term effects of patients weaning off PFC liquid ventilation remain largely unknown and are not tested in this thesis. Moreover, PFC is an umbrella term for PFD and PFOB used in this thesis; it is noteworthy that there is an incomplete investigation of both PFC variants across all experimental modules due to supply problems with PFD. However, including PFOB is sufficient as liquid ventilation clinical trials exclusively use PFOB for its imaging contrast property, and PFD is more used in animal studies.

#### Lack of lung microenvironment and limited cell types

Immune cells were not included in this AEC-exclusive. The airway epithelium functions through the interaction of immune cells, such as resident macrophages, that are localised in alveolar spaces. This limits the ability to fully assess the immune modulation of AEC in response to PFC exposure. Additionally, no primary alveolar cell culture or similar cell type was used to compare outcomes with PFC-exposed A549 culture. Considering the scarcity of human primary alveolar AEC, it may be worth resorting to immortalised cell line with surfactant expression to assess the effects of PFC on alveoli with or without surfactant, as there have been mixed suggestions as to whether PFC can be a sole replacement of surfactant during liquid ventilation or co-administrations are needed.

#### Interpretation of Autophagy data

Autophagy is a dynamic process, and autophagic flux refers to completing this process [50]. SEQ and LC3BII are degraded at the end of flux, and Western blot results of these two proteins must be interpreted cautiously. For instance, LC3BII blot results reflect the number of autophagosomes at a given time point and do not reflect the true rate/frequency of flux [58]. An increase in flux may result in the degradation of autophagy markers and appear as faint signals on Western blot. This thesis uses SEQ and LC3BII abundance to indicate autophagy during the experiment endpoint. To accurately measure flux, inhibition of lysosome function, such as through the administration of bafilomycin, will accumulate undegraded LC3BII signals in the sample, and the accumulated signals compared to untreated samples will represent the level of flux. Additionally, SEQ is highly transcriptionally regulated; polymerase chain reaction (PCR) analysis of SEQ transcripts may provide additional clues to autophagy modulation after PFC exposure.

#### Future Directions

To address the limitations of PFC exposure mentioned, a microfluidic device of organ-on-a-chip technology presents a powerful platform with several advantages. Firstly, fluid flow and shear stress are precisely controlled in these devices [59], allowing for recapitulation of the dynamic flow of PFC during liquid ventilation and cellular behaviour study under physiologically relevant conditions. Secondly, more cell types can be incorporated into this system to create a complex, realistic airway-on-a-chip. Performing PFC exposure in this system allows in-vitro modelling of the respiratory system instead of only the epithelial aspect.

Another avenue for future research involved exploring different formulations of PFC emulsions or PFC compounds. An emulsion is a mixture of immiscible liquids; PFC emulsions have been used as experimental artificial oxygen carriers in blood for their exceptional oxygen-carrying capability [60]. The diverse chemical properties of various formulations may enhance biocompatibility with optimised physical properties for liquid ventilation. Investigating various PFC formulations could help identify the optimal PFC for use in different populations with the same condition, such as adult and neonatal respiratory distress syndrome. Drug delivery via PFC liquid ventilation is also emerging as a promising avenue for targeted respiratory therapies. PFCs serve as great carriers for drugs, considering their low surface tension property that allows effective coverage of the lung surface and delivers drugs directly to the AEC surfaces. We are making emulsions containing azithromycin, and are overcoming issues with cell toxicity using this method (Supp Fig 1). Developing this approach may increase drug efficacy for improved clinical outcomes. However, in-vitro studies are necessary, considering PFC emulsions or drug emulsions will be utilised for optimised drug delivery and asses any potential toxicities before in vivo studies.

Furthermore, the ability to culture primary cells using the model employed in this thesis means PFC-based therapies can be optimised through personalised approaches. By considering the innate biological variations between patients and respective conditions, tailored liquid ventilation strategies will increase the technique’s safety and efficacy while opening the possibility of liquid ventilation for other unexplored respiratory conditions.

## Conclusion

In summary, this thesis demonstrates the practicality of an air-liquid interface organotypic model to investigate the interactions between PFC and human airway epithelium for the medical implications of liquid ventilation. Findings from 16HBE14o-differentiated hSABCi-NS1.1 and primary nAEC grown at air-liquid interface demonstrated PFCs’ inert nature and supported PFC’s biocompatibility property established from animal and clinical studies. Specifically, PFC exposure apically at the condition tested did not induce significant apoptosis, evident from minimal PARP cleavage, and autophagy remained unmodulated, evident from basal level SQSTM1 and LC3B abundance in organotypic culture. Also, hSABCi and nAEC preserve barrier function with stable TEER and tight junction protein expression during exposure. The exposure conditions in this thesis were intentionally kept straightforward to provide foundation work for future in-vitro investigations of PFC. Future investigations should build upon the 3D organotypic architecture employed in this thesis. Introduce conditions that mimic PFC medical applications in vivo to fully realise the potential of PFC beyond just an alternative of respiratory care towards innovative solutions for human endeavours in low-oxygen environments.

## Acknowledgements

Thank you to Miss Izabella Sabatini and Dr Dishant Vikram Bhai Patel for technical support.

## Author contributions

ER was the research founder, conducted the analyses, performed laboratory experiments, co-drafted the manuscript and served as corresponding author. SWHL conducted the analyses, performed laboratory experiments and co-drafted the manuscript. FOC conducted the analyses, performed laboratory experiments and co-drafted the manuscript. RDQ conducted the analyses and co-drafted the manuscript. Each author contributed intellectual effort, and read and approved the final manuscript.

---

**Supp Fig 1 Figure 4:**
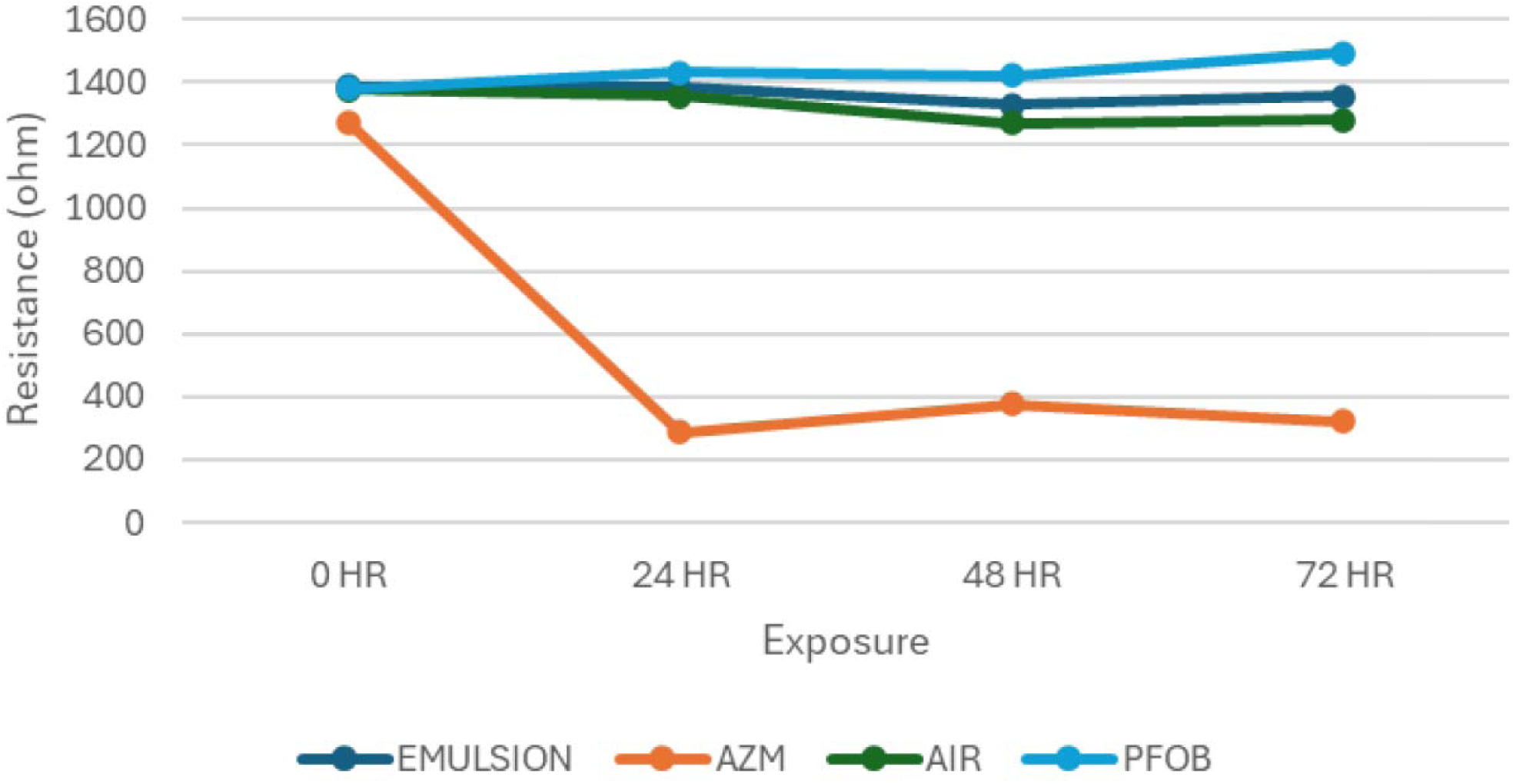
TEER measurements of epithelial monolayers treated with various PFC formulations over time. Transepithelial electrical resistance (TEER) values were measured at 0-, 24-, 48-, and 72-hours following treatment with the following conditions: (1) perfluorocarbon (PFC) emulsion, (2) azithromycin-loaded PFC emulsion (AZM-PFC emulsion), (3) air, and (4) PFOB. TEER values are expressed as ohms of baseline resistance (0 h), indicating barrier integrity. Data are presented as mean ± SD from n = 3 independent replicates. A decrease in TEER over time indicates compromised epithelial barrier function, while stable or increasing values suggest intact or improving barrier properties.

